# Intra-hypothalamic circuit orchestrates β-endorphin release following coital ejaculation in male mice

**DOI:** 10.1101/2024.09.18.613624

**Authors:** Xi Zha, Zhuo-Lei Jiao, Shuai-Shuai Li, Xiao-Yao Liu, Xing-Yu Li, Yi-Zhuo Sun, Xiao-Jing Ding, Meng-Tong Gao, Shu-Chen Gao, Ai-Xiao Chen, Jun-Kai Lin, Wen Zhang, Xuan-Zi Cao, Yan-Li Zhang, Rong-Rong Yang, Chun Xu, Xiao-Hong Xu

## Abstract

Survey-based evidence suggests that men experience a distinct post-ejaculation affective state^1,2^, marked by intense pleasure sometimes compared to the euphoric rush from intravenous injection of opioid drugs such as heroin^3^. However, the intrinsic neural circuit mechanisms underlying the ejaculation-triggered affective state remain unclear. Here, we discovered that *Calbindin1*-expressing (*Calb1+*) neurons in the preoptic area (POA) of the hypothalamus, an evolutionarily conserved regulatory region for male mating behavior^4^, are specifically activated during ejaculation in male mice. Inhibiting POA *Calb1+* neurons prolongs mating and delays ejaculation. Importantly, POA *Calb1+* neurons transmit the ejaculation signal and activate proopiomelanocortin-expressing (*Pomc+*) neurons in the arcuate nucleus of the hypothalamus, which show robust and sustained activity lasting for tens of seconds, specifically upon ejaculation. This activity is accompanied by elevated levels of β-endorphins^5^, opioid peptides secreted by *Pomc+* neurons, post-ejaculation in male mice. Optogenetic activation of *Pomc+* neurons increases β-endorphins levels and conditioned placed preference, similar to ejaculation. Conversely, intracerebroventricular (i.c.v.) infusion of drugs blocking *Pomc* neuropeptides signaling eliminates ejaculation-conditioned place preference. Collectively, these results elucidate an intra-hypothalamic circuit from POA *Calb1+* neurons to arcuate *Pomc+* neurons that coordinate β-endorphin release with ejaculation, shedding light on the neurobiological basis of the post-ejaculation affective state.

## Main Text

Extensive research in rodents has mapped a neural circuit spanning from sensory input to hypothalamic and midbrain regions that drives the stereotyped and conserved male mating behaviors observed across mammalian species^4,6,7^. Central to this male mating circuit is the preoptic area (POA) of the hypothalamus, which plays a critical, evolutionarily conserved role in regulating male mating. Recording and functional manipulation experiments in mice have shown that specific populations of POA neurons expressing estrogen receptor 1 (*Esr1*) and tachykinin receptor 1 (*Tacr1*) are essential for initiating mounting and intromission (also known as pelvic thrusts)^8,9^. These mating-promoting POA neurons project to the ventral tegmental area (VTA), where they activate dopamine neurons^10^, leading to dopamine release in the nucleus accumbens (NAc) during mating^9,11^. The release of another neuromodulator, oxytocin, was also detected during mount and intromission^12^.

Ejaculation, a complex physiological process involving the coordination of various muscles to release semen^13,14^, succeeds the acts of mounting and intromission. Ejaculation marks the conclusion of the consummatory phase of mating, and is commonly associated with a profound sense of pleasure, as reported by men in surveys^1,13,15^. Previous studies in rats have identified neurons in the spinal cord necessary for coordinating the reflexive motor and autonomic outputs associated with ejaculation^16^. Although neurons in the brain are thought to regulate the timing of ejaculation through interactions with the spinal cord “ejaculation generator” neurons^17,18^, the molecular identity of supraspinal neurons that regulate ejaculation remains poorly understood.

Noteworthily, the mental state accompanying ejaculation is distinct from that experienced during mount and intromission. This distinction indicates that while dopamine and oxytocin are released during ejaculation as in mount and intromission^11,12,14^, other neural pathways and neurochemicals are likely to be specifically recruited during ejaculation to mediate the post-ejaculation affective state. Intriguingly, the euphoric sensation triggered by intravenous heroin injection is sometimes compared to the pleasure derived from ejaculation^3,19^, suggesting the possible involvement of the endogenous opioid system in the post-ejaculation affective state.

While recent studies have begun to uncover activity changes in molecularly defined neuronal populations related to post-ejaculation mating suppression^20,21^, the neural mechanisms underlying the unique affective state following ejaculation remain largely underexplored^18^. We hypothesize that identifying neurons specifically activated during ejaculation in the central nervous system could shed light on the neural processes underlying the post-coital affective state. Classical studies utilizing immunostaining of immediate early genes (IEGs), such as c-Fos, as markers of neuronal activation have shown that more POA neurons are labeled when male rodents achieve ejaculation compared to intromission alone^22–24^. Therefore, we began by investigating the identity of POA neurons that are selectively activated during ejaculation in male mice.

### Act-seq reveals selective activation of POA Calb1+ neurons during ejaculation

We dissected preoptic area (POA) tissues from C57BL/6 male mice approximately 30 minutes after they displayed intromission or ejaculation during a standard mating test with a female (**Fig. 1a**). The total duration of the behavioral assay was matched between the two groups to ensure comparability. Tissues from three animals per condition were pooled and processed with Act-D to inhibit transcription induced by tissue dissociation before library construction^25,26^. In total, 27,447 sequenced nuclei (with 15,453 from the intromission group and 11,768 from the ejaculation group) were identified as neurons transcriptionally and processed for further analysis.

**Figure 1.**
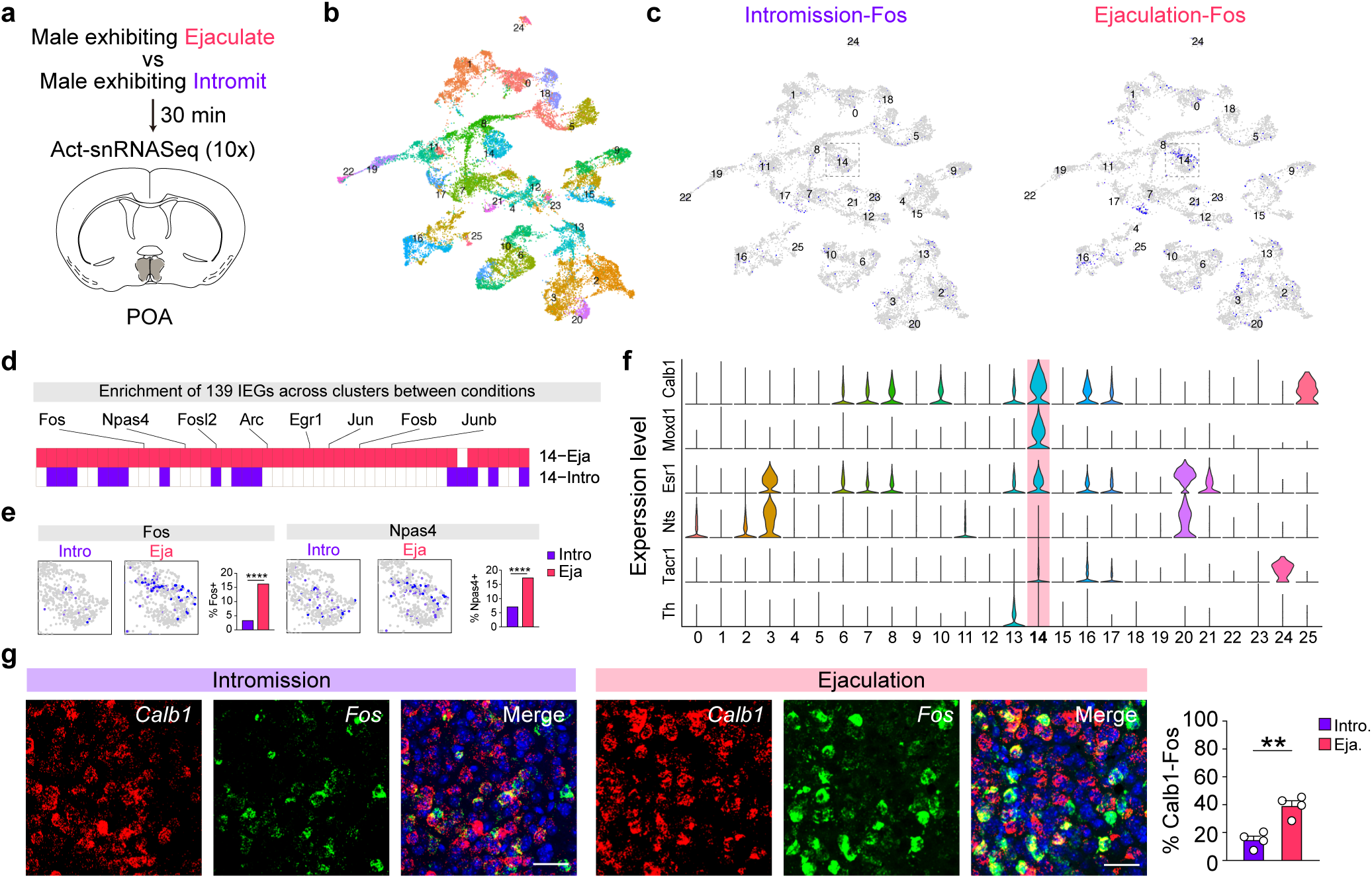
Single-nucleus RNA-seq reveals selective activation of POA *Calb1+* neurons during ejaculation. **a.** Schematic illustration of the preoptic area (POA, gray) dissected from a coronal brain section of C57BL/6 male mice sacrificed 30 minutes after mating tests. Males either achieved intromission (blue) or ejaculation (red) during the test. Three males were included in either group. **b.** Single-nucleus RNA-seq analysis identifies 26 neuronal clusters in the POA. **c.** Distribution of *Fos+* signals across clusters in cells and clusters from the intromission (left) or ejaculation (right) condition. **d.** Illustrating the immediate early genes (IEGs) significantly enriched in Cluster #14 under the ejaculation condition (top, red) and the intromission condition (bottom, blue). A larger number of IEGs showed significant enrichment in Cluster #14 during ejaculation condition compared to intromission condition. The full list of 139 IEGs is provided in Extended Data Fig. 2a. **e.** A zoom-in view of cluster #14 with the quantification in the bar graph showing a significantly higher percentage of cluster #14 neurons expressing *Fos* (left) and *Npas4* (right) in the ejaculation condition compared to the intromission condition. **f.** Relative expression levels of marker genes (shown on the left) across the 26 neuronal clusters. Cluster #14 is characterized as *Calb1+/Moxd1+/Esr1+*. **g.** Examples RNAscope (fluorescent *in situ* hybridization) images on the left and quantification on the right, showing that more *Calb1+* neurons co-expressing *Fos* in POA after males achieving ejaculation (red) than intromission alone (blue). N = 4 males/either group. Scale bar, 50 μm. Values are presented as mean ± SEM. ** p < 0.01, ****p < 0.0001.

Analyzing the snRNA-seq data of these 27,447 nuclei generated 26 POA neuronal clusters (**Fig. 1b**), with each cluster consisting of comparable percentage of cells dissociated from the intromission or ejaculation conditions (**Fig. 1c**, Extended Data Fig.1a-1b), suggesting minimum tissue dissection variations between the two conditions. When examining the POA as a whole, we found more robust activation of POA neurons during ejaculation compared to intromission, as indicated by significant upregulation of many immediate early genes (IEGs) known to be sensitive to neuron activity in the ejaculation condition (Extended Data Fig. 1c), consistent with previous results^22–24^.

Further analysis of the enrichment ratio of IEGs across the 26 POA neuron clusters revealed activation of distinct neuron clusters during ejaculation and intromission (**Fig 1d**, Extended Data Fig. 2a). Some clusters, including clusters #3, #16, and #24, exhibited robust activation both during intromission and ejaculation (Extended Data Fig. 2a). Intuitively, this may suggest accumulative activation of these neuronal clusters leading up to ejaculation. In fact, these three clusters expressed *Esr1* or *Tacr1* (Extended Data Fig. 2b) as the molecular markers, aligning well with previously identified POA neuron populations known to be involved in male mating^8,9^.

In contrast, clusters 14 and 7 were primarily activated in the ejaculation but not the intromission condition (Extended Data Fig. 2a). Intriguingly, cluster #14, which showed the most pronounced activation during ejaculation (**Fig.1d-1e,** Extended Data Fig. 2a), was characterized by the expression of *Calb1* and *Moxd1* (**Fig. 1f**), both of which are markers of the sexually dimorphic nucleus of the preoptic area (SDN-POA)^27,28^. Despite extensive histological characterization and classical lesion studies, the role of SDN-POA in male mating behavior remained unclear^29–31^. Notably, *Calb1* is also enriched in cluster #16, an excitatory neuron cluster activated during both intromission and ejaculation and also characterized by the expression of *Adcyap1*(Extended Data Fig. 2a-2b). Together, these results support activation of POA *Calb1+* neurons during ejaculation.

To validate this finding, we conducted fluorescent *in situ* hybridization (RNAscope) for *Fos*, along with *Calb1* and *Adcyap1*, on brain sections from males that had either achieved intromission or ejaculation in a mating test (Extended Data Fig. 3a). We found a significantly higher number of *Calb1+* neurons being labeled with *Fos* after males achieved ejaculation compared to intromission alone (**Fig. 1g**). Furthermore, ∼ 80% of *Adcyap1* neurons co-expressed *Calb1*, and these neurons also showed increased *Fos* expression following ejaculation (Extended Data Fig. 3b). Collectively, these findings identify subsets of POA *Calb1+* neurons as neurons specifically activated during ejaculation.

### POA Calb1+ neuron activity facilitates ejaculation

To investigate the temporal profile of POA *Calb1+* neuron activity during male mating, we monitored Ca^2+^ transients in these neurons (**Fig. 2a**). We injected adeno-associated viruses (AAVs) encoding Cre-dependent GCaMP6s (Cre-on strategy) into the POA of *Calb1-Cre* male mice (**Fig. 2b**) and implanted an optic fiber. For comparison, we injected AAVs encoding Cre-deleted GCaMP6s (Cre-off strategy) into the same region in other *Calb1-Cre* male mice to label *non-Calb1* neurons (**Fig. 2b**) and performed similar experiments. Immunostaining demonstrated co-labeling of GCaMP6s with Calb1 in males injected with Cre-on AAVs, while little co-labeling was observed in males injected with Cre-off virus (**Fig. 2c**), confirming the specificity of the viral targeting strategy.

**Figure 2.**
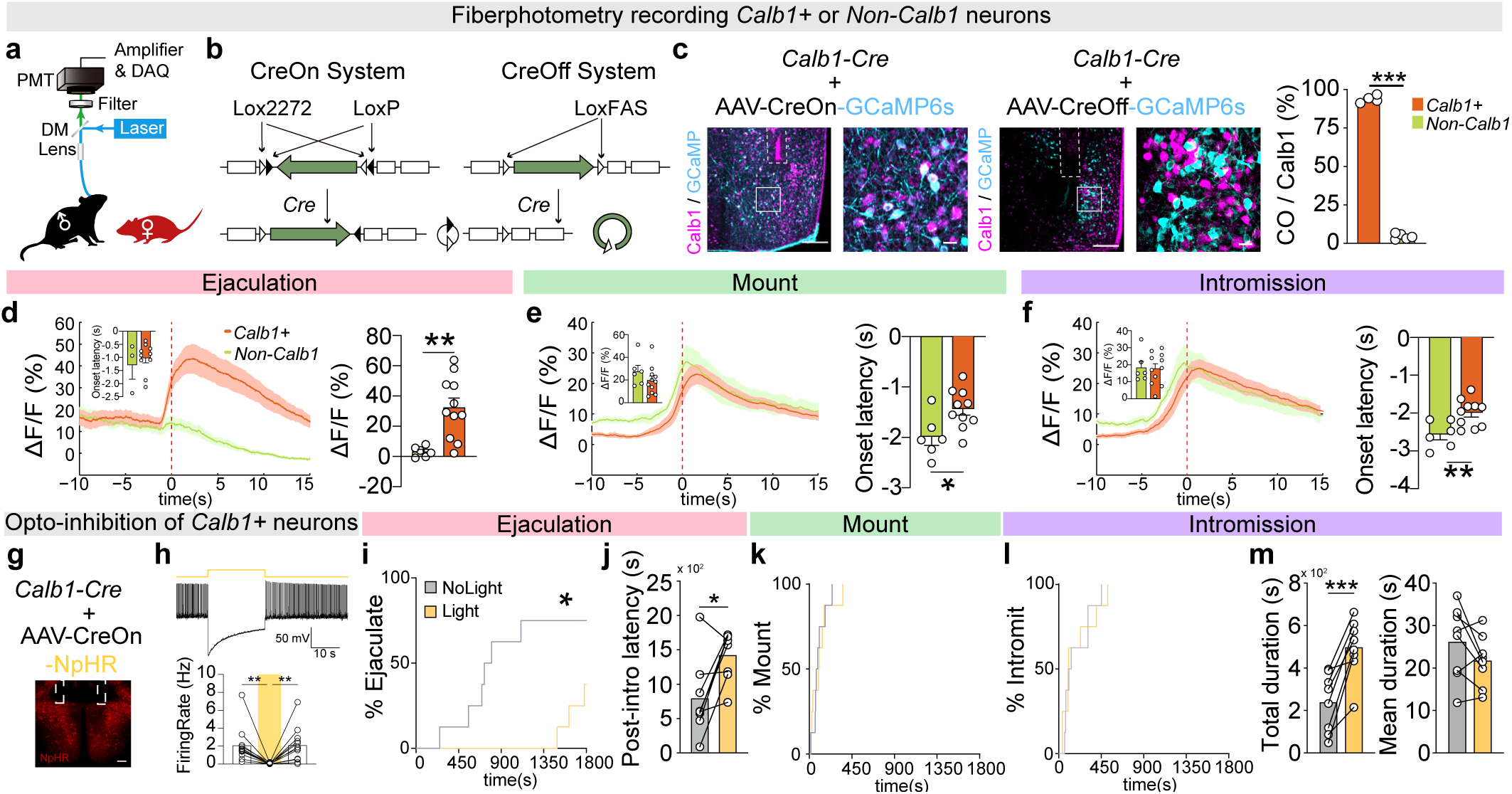
Activity of *Calb1+* neurons facilitates the transition from intromission to ejaculation during male mating. **a.-b.** Experimental setup (**a**) and the details of the Cre-On and Cre-Off viral strategy (**b**) to label and record the activity of POA *Calb1+* and *non-Calb1* neurons. **c.** Representative images of *Calb1+* and *non-Calb1* neurons following injection of AAVs encoding Cre-On or Cre-off GCamp6s into the POA of *Calb1-Cre* male mice are shown on the left, with the images on the right highlighting the area outlined by the white box on the left. Scale bar, 200 μm (left); 20 μm (right). The quantification is shown on the (right). n = 4 males injected with Cre-On virus and 5 males injected with Cre-Off males. **d.-f.** Average changes in GCamp6s signals (ΔF/F) recorded from *Calb1+* (orange) or *non-Calb1* neurons (light green) around ejaculation (**d**), mounting (**e**), and intromission (**f**). Behavior-triggered averages are aligned to the onset of each behavior, marked as the time “0”. Quantification of latency and average ΔF/F signals is presented in the accompanying bar graphs. n = 11 males injected with Cre-On virus and 6 males injected with Cre-Off males. **g.** *Calb1-Cre* male mice were injected with AAVs encoding Cre-On NpHR. A representative image at the bottom shows NpHR expression in the POA, with dashed lines indicating implanted optic fibers. Scale bar, 200 μm. **h.** A representative raw trace on the top and the quantification at the bottom show that light delivery (indicated by the yellow bar) in the brain slices significantly reduced the firing rates of *Calb1+* neurons expressing NpHRs. n = 12 cells recorded from 2 mice. **i.-m.** Optogenetic inhibition of *Calb1+* neurons reduced the percentage of males that ejaculated following female introduction at time ‘0’ (**i**), increased ejaculation latency following the 1^st^ intromission (post-intro) (**j**), without affecting the percentage of males that mounted (**k**) or intromitted (**l**) following female introduction at time “0”. Additionally, it extended the total duration of intromission without altering the mean duration of each intromission bout (**m**). n = 8 males. Values are presented as mean ± SEM. *p < 0.05, **p < 0.01, ***p < 0.001.

Clearly, we found a significant increase in GCaMP6s signals during ejaculation in *Calb1+* but not *non-Calb1* neurons, supporting ejaculation-specific activation of POA *Calb1+* neurons (**Fig. 2d**, Extended Data Video 1-2), consistent with the Act-seq results. Notably, the activity in POA *Calb1+* neurons preceded ejaculation onset (**Fig. 2d)**, which was identified by the characteristic stiffening of posture observed in males during the final intromission^32^ (Extended Data Video 1-2), and last for over 15 seconds (signal latency, -1.0 ± 0.2 s; duration, 16.3 ± 1.9 s, n = 10). Importantly, in *Calb1-Cre* male mice injected with the control EYFP viruses, no changes in fluorescent signals were detected at any stage of mating (Extended Data Fig. 4), suggesting that the GCaMP6s signals detected in *Calb1+* neurons during ejaculation were unlikely caused by motion artifacts.

Increased GCaMP6s signals were also observed in both *Calb1+* and *non-Calb1* neurons during mount and intromission, with the signals in both populations reaching similar levels (**Fig. 2e-2f**). However, the GCaMP6s signals in *Calb1+* neurons significantly lagged behind those of *non-Calb1* neurons by approximately 0.5 seconds during mount and intromission (**Fig. 2e-2f**). This temporal difference suggests possible distinct roles played by *Calb1+* and *non-Calb1* neurons during male mating. To test this, we expressed the light-sensitive cation channel Channelrhodopsin 2 (ChR2) in POA *Calb1+* or *non-Calb1* neurons using the genetic strategy described above and then optogenetically activated these neurons (Extended Data Fig. 5a-5c), we found that while stimulating *non-Calb1* neurons promoted mounting behavior (Extended Data Fig. 5d-5f), stimulating *Calb1+* neurons not only failed to promote mounting but also appeared to inhibit it (Extended Data Fig. 5g-5i).

Next, we optogenetically inhibited POA *Calb1+* neurons during male mating. AAVs encoding *Cre*-dependent halorhodopsin (NpHR) were injected bilaterally into the POA of *Calb1-Cre* males, followed by the implantation of bilateral optic fibers (**Fig. 2g**). Continuous yellow light (590 nm), shown to reversibly suppress the activity of NpHR-expressing *Calb1+* neurons in brain slice recordings (**Fig. 2h**), was delivered when the males were within one body length (∼ 9 cm) of the female during mating. Remarkably, we found that optogenetic inhibition of POA *Calb1+* neurons significantly delayed ejaculation (**Fig. 2i-2j**) but had little effects on the initiation of mount and intromission behavior (**Fig. 2k-2l**). This led to an increased total duration of intromissions before ejaculation, while the average duration of each intromission remained unaffected during optogenetic inhibition of POA *Calb1+* neurons (**Fig. 2m**). This sharply contrasted with optogenetic inhibition of POA *Esr1+* neurons, which largely blocked mounting behavior^8^. These results highlight the distinct role played by POA *Calb1+* neurons in facilitating the progression from intromission to ejaculation during male mating.

Importantly, similar effects of delayed ejaculation were also observed when AAVs encoding Cre-dependent Caspase 3 were injected into the POA of *Calb-Cre* males to permanently ablate POA *Calb1+* neurons (Extended Data Fig. 6a-6b). These ablated males were comparable to control males in their ability to initiate mounting and intromission (Extended Data Fig. 6c-6d) but experienced difficulty achieving ejaculation, as indicated by increased ejaculation latency (Extended Data Fig. 6e-6f), with a prolonged total duration of intromission compared to those injected with a control virus (Extended Data Fig. 6d). Together, these findings demonstrate that POA *Calb1+* neurons play a critical and specific role in facilitating ejaculation.

### Arcuate Pomc+ neurons are downstream of POA Calb1+ neurons

The ejaculation-specific activation of POA *Calb1+* neurons provided an entry point to trace the downstream neuronal substrates that may underlie the post-ejaculation affective state. To explore this, we compared the axon projection of POA *Calb1+* and *non-Calb1* neurons by analyzing the mCherry signals in *Calb1-Cre* male mice injected with either Cre-on or Cre-off ChR2-mCherry viruses (Extended Data Fig.7). We observed significantly denser innervation of the arcuate nucleus (ARC) and the adjacent ventromedial hypothalamus (VMH) for POA *Calb1+* neurons compared to *non-Calb1* neurons (**Fig. 3a-3b**, Extended Data Fig. 7b-7c).

**Figure 3.**
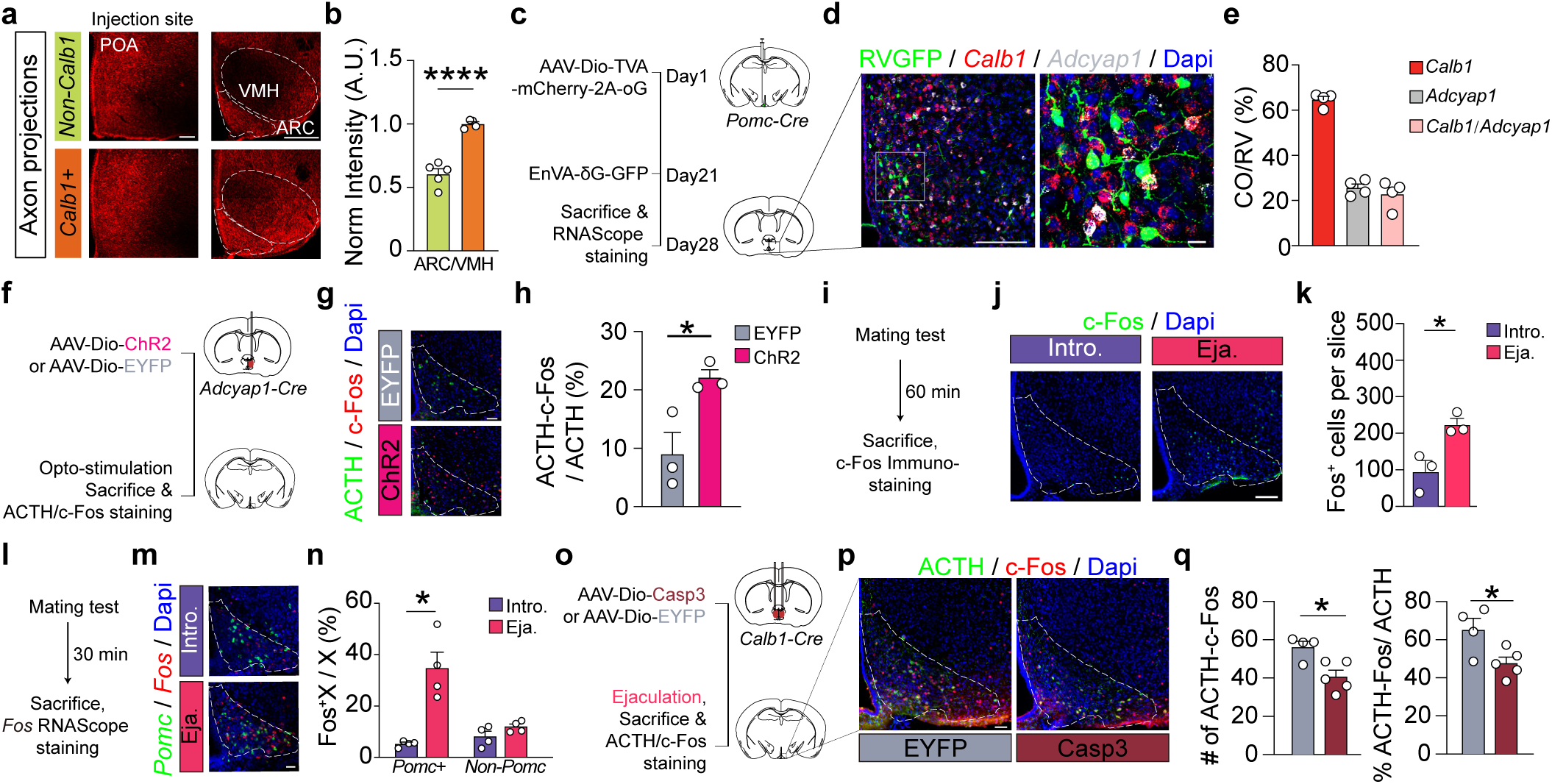
Arcuate *Pomc+* neurons are downstream of POA *Calb1+* neurons and are activated post-ejaculation. **a.-b.** AAVs encoding Cre-Off or Cre-On ChR2-mCherry were injected unilaterally into the POA of *Calb1-Cre* male mice to label *non-Calb1* and *Calb1+* neurons, respectively. Representative images in (**a**) show mCherry signals in POA (left) and in arcuate nucleus (ARC, outlined) and the ventromedial hypothalamic nucleus (VMH, outlined) on the right. Scale bar, 200 μm. Quantification in (**b**) shows denser innervation of ARC/VMH area by *Calb1+* neurons. n = 4 males for the *Calb1+* group and 5 males for *non-Calb1* group. **c.-e.** Pseudo-rabies-virus-mediated retrograde tracing from ARC *Pomc+* neurons identifies inputs from POA *Calb1+* neurons, including a subset of *Calb1+* neurons co-expressing *Adcyap1*. The viral retrograde tracing strategy is shown in (**c**); representative imaging showing co-labeling of RV-GFP, *Calb1*, and *Adcyap1* in POA (**d,** scale bar, 200 μm for the image on the left and 20 μm for the image on the right), with quantification in (**e**). n = 4. **f.-h.** Optogenetic activation of POA *Adcyap1+* neurons increases the percentage of ARC *Pomc+* neurons, identified by ACTH immunohistochemistry, co-expressing c-Fos. The experimental procedure is depicted in (**f**), representative images in ARC (outlined) showing ACTH and c-Fos co-localization in (**g,** scale bar, 50 μm), and quantification is presented in (**h**). n = 3 ChR2 and 3 EYFP males. **i.-k.** Increased c-Fos expression in ARC following ejaculation (Eja.) compared to intromission (Intro.) alone. The experimental procedure is shown in (**i**), the representative images of ARC (outlined) c-Fos expression are shown in (**j,** scale bar, 100 μm) with the quantification shown in (**k**). n = 3 intromission and 3 ejaculation males. **l.-n.** Ejaculation (Eja.) leads to increased percentage of neurons co-expressing *Fos* in ARC *Pomc+* but not *non-Pomc* population. The experimental procedure is shown in (**l**), the representative images of *Fos* and *Pomc* double fluorescent *in situ* in ARC (outlined) are shown in (**m,** scale bar, 50 μm) with the quantification shown in (**n**). n = 4 intromission and 4 ejaculation males. **o.-q.** Selective ablation of POA *Calb1+* neurons leads to reduced post-ejaculaiton c-Fos expression in ARC *Pomc+* neurons, identified by ACTH immunohistochemistry. The viral strategy and experimental procedure is shown in (**o**), representative images of c-Fos expression in ARC (outlined) in (**p**, scale bar, 50 μm) and quantification in (**q**). n = 5 caspase and 4 EYFP males. Values are presented as mean ± SEM. *p < 0.05, ****p < 0.0001.

Given that proopiomelanocortin (*Pomc*)-expressing neurons, known to regulate satiety and serve as a source of endogenous opioid endorphins^33^, reside in the ARC/VMH region, we investigated whether *Pomc+* neurons are downstream of POA *Calb1+* neurons. Our pseudo-rabies retrograde tracing in *Pomc-Cre* males (**Fig. 3c**, Extended Data Fig. 8a) revealed intense labeling of POA neurons upstream of arcuate *Pomc+* neurons, with approximately 70% co-expressing *Calb1* (**Fig. 3d-3e**), confirming that POA *Calb1+* neurons innervate *Pomc+* neurons. Additionally, the subset of POA *Calb1+* neurons co-expressing *Adcyap1*, a subtype of POA glutamatergic neurons, were also retrogradely labeled (**Fig. 3e**), providing a neural pathway through which activation of POA *Calb1+/Adcyap1+* neurons could directly excite arcuate *Pomc+* neurons. Supporting this, optogenetic activation of POA *Adcyap1+* neurons led to increased c-Fos expression in arcuate *Pomc+* neurons (**Fig. 3f-3h**).

Moreover, in males that achieved ejaculation a higher number of c-Fos+ signals was detected in the arcuate nucleus compared to males who only experienced intromission (**Fig. 3i-3k**). Further analysis revealed that approximately 40% of *Pomc*+ neurons co-expressed *Fos* in males post-ejaculation, compared to only about 5% in males that experienced intromission only (**Fig. 3l-3n**). Notably, the percentage of nearby *non-Pomc* neurons expressing *Fos* did not significantly increase following ejaculation (**Fig. 3n**), highlighting the specific activation of *Pomc+* neurons during this process. Additionally, ablating POA *Calb1+* neurons significantly reduced c-Fos*+* signals in *Pomc+* neurons after ejaculation (**Fig. 3o-3q**). These findings collectively establish arcuate *Pomc+* neurons as a key downstream target of POA *Calb1+* neurons that is activated post-ejaculation.

### Sustained Pomc+ neuron activity during ejaculation

To further explore the temporal dynamics of *Pomc+* neuron activity, we injected AAVs encoding Cre-dependent GCaMP6s into the arcuate nucleus of *Pomc-Cre* male mice and recorded Ca^2+^ transients from these neurons during male mating through the implanted optic fiber (**Fig. 4a**). We found specific activation of arcuate *Pomc+* neurons exclusively during ejaculation, as evidenced by a pronounced surge of GCaMP6s signals during ejaculation, with negligible activity observed throughout other mating behaviors or in EYFP-expressing control animals (**Fig. 4b-4c**, Extended Data Video 3, Extended Data Fig. 4e-4f). The GCaMP6s signals in *Pomc+* neurons were time-locked to the moment of ejaculation and emerged later than the signals in POA *Calb1+* neurons (**Fig. 4b**, Extended Data Fig.8b), consistent with *Pomc+* neurons being downstream of POA *Calb1+* neurons. Remarkably, the ejaculation-associated activity in *Pomc+* neurons lasted for an average of 50 seconds before subsiding, significantly outlasting the activity observed in POA *Calb1+* neurons during ejaculation (**Fig. 4b**, Extended Data Fig.8b). In stark contrast, adjacent neurons expressing agouti-related neuropeptide (*Agrp*) showed no such elevated activity (**Fig. 4d-4f**), demonstrating that ejaculation-related activity is specific to *Pomc+* neurons.

**Figure 4.**
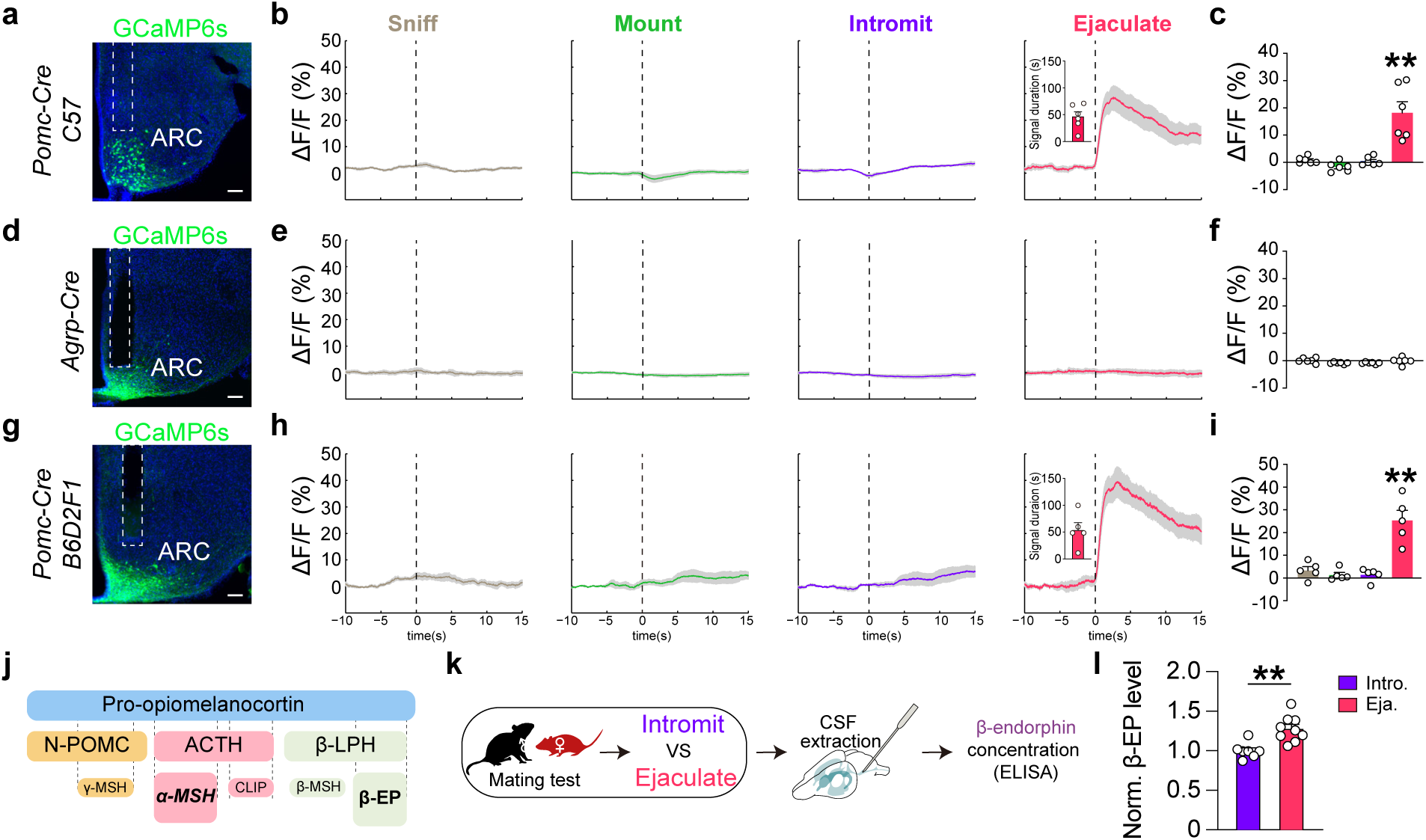
Sustained activation of ARC *Pomc+* neurons and elevated CSF β-endorphin levels post-ejaculation. **a, d, g.** Representative images showing expression of GCamp6s in ARC *Pomc+* (**a, g**) and *Agrp+* (**d**) neurons after injection of AAVs encoding Cre-On GCamp6s into the *Pomc-Cre* or *Agrp*-*Cre* male mice of C57BL/6 background (**a, d**), or *Pomc-Cre* male mice of the B6D2F1 hybrid background (**g**), for fiber photometry recording experiments. Dashed lines depicting implanted optical fibers. Scale bar, 100 μm. **b, e, h.** Behavior-triggered averages of ΔF/F signals during the indicated behaviors (columns) for the recorded population (rows), with the corresponding signal duration values for ejaculation shown in the insets for *Pomc+* neurons. Quantification of signal intensity is displayed on the right in **c, f, i.** n = 6 males for C57BL6/*Pomc+* group, 6 males for C57BL6/*Agrp+* group, 5 males for B6D2F1/*Pomc+* group. **j.-k.** Schematics showing the processing of POMC and the resulting mature neuropeptides in (**j**) and the experimental procedure for extracting CSF to measure β-endorphin levels after male mating. **l.** β-endorphin (β-EP) levels normalized (norm.) to the mean value of the intromission (intro.) group show elevation in ejaculated (Eja.) males. n = 7 intromission and 9 ejaculation males. Values are presented as mean ± SEM. **p < 0.01.

Furthermore, F1 hybrid male offsprings from crosses between DBA/2 males and C57BL/6 females show a faster recovery to mating after ejaculation compared to the parental C57BL/6 line^34^. However, in such F1 hybrid males (Extended Data Fig.8c), a similar pattern of sustained activation in *Pomc+* neurons was observed during ejaculation, mirroring that seen in C57BL/6 males, (**Fig. 4g-4i**, Extended Data Video 4, GCaMP6s signal duration: C57, 46.5 ± 9.3s, n = 6, F1 hybrid, 53.5 ± 14.2s, n = 5). These findings suggest that *Pomc+* neurons in male mice exhibit prolonged activation specifically associated with ejaculation, regardless of genetic background or variations in post-ejaculation mating suppression.

*Pomc+* neurons are known to secrete at least two distinct types of neuropeptides: melanocortin peptides, including α, β, γ-MSH, ACTH, and opioids like β-endorphin, that are derived from precursors encoded by the *Pomc* gene^5^ (**Fig 4j**). Among them, β-endorphin is secreted by *Pomc+* neurons into the ventricle and is particularly noteworthy for its rewarding effects^35^. To investigate whether β-endorphin is released following ejaculation, we collected cerebrospinal fluid (CSF) from males immediately after ejaculation with males that experienced only intromission as controls (**Fig. 4k**). Our ELISA measurements revealed a significant elevation in CSF β-endorphin levels following ejaculation (**Fig. 4l**). This evidence underscores the ejaculation-specific activation of arcuate *Pomc+* neurons and the subsequent release of β-endorphin, likely along with other neuropeptides, following ejaculation, suggesting their possible role in regulating the post-ejaculation emotional state.

### Regulation of post-ejaculation affective state by Pomc+ neurons

To evaluate the affective state following ejaculation in male mice, we utilized the conditioned place preference (CPP) test ^36,37^. Male mice were allowed to interact with a female in their home cage until ejaculation occurred or a similar amount of time had passed. Immediately afterward, they were placed in one of two distinct chambers with unique textures within a three-chambered apparatus, in a roughly counter-balanced manner (**Fig. 5a**). Following standard procedures for studying reward stimuli^36,37^, the ejaculation condition was paired with the initially less preferred chamber on the pre-test (day 1). On the test day after the conditioning phase (day 4), the males were allowed to explore the entire apparatus, and the time they spent in each chamber was recorded (**Fig. 5a**). This post-conditioning data was then compared to the pre-test measurements to assess the changes in time spent in and the preference scores of the paired chambers (**Fig. 5b**). Our results demonstrated that, after conditioning, wildtype males of both the C57BL/6 and the B6D2F1 hybrid background spent significantly more time in the chamber paired with ejaculation (**Fig. 5c**). This shift increased the preference score for the ejaculation-paired chamber (**Fig. 5c**), demonstrating the rewarding effect of ejaculation.

**Figure 5.**
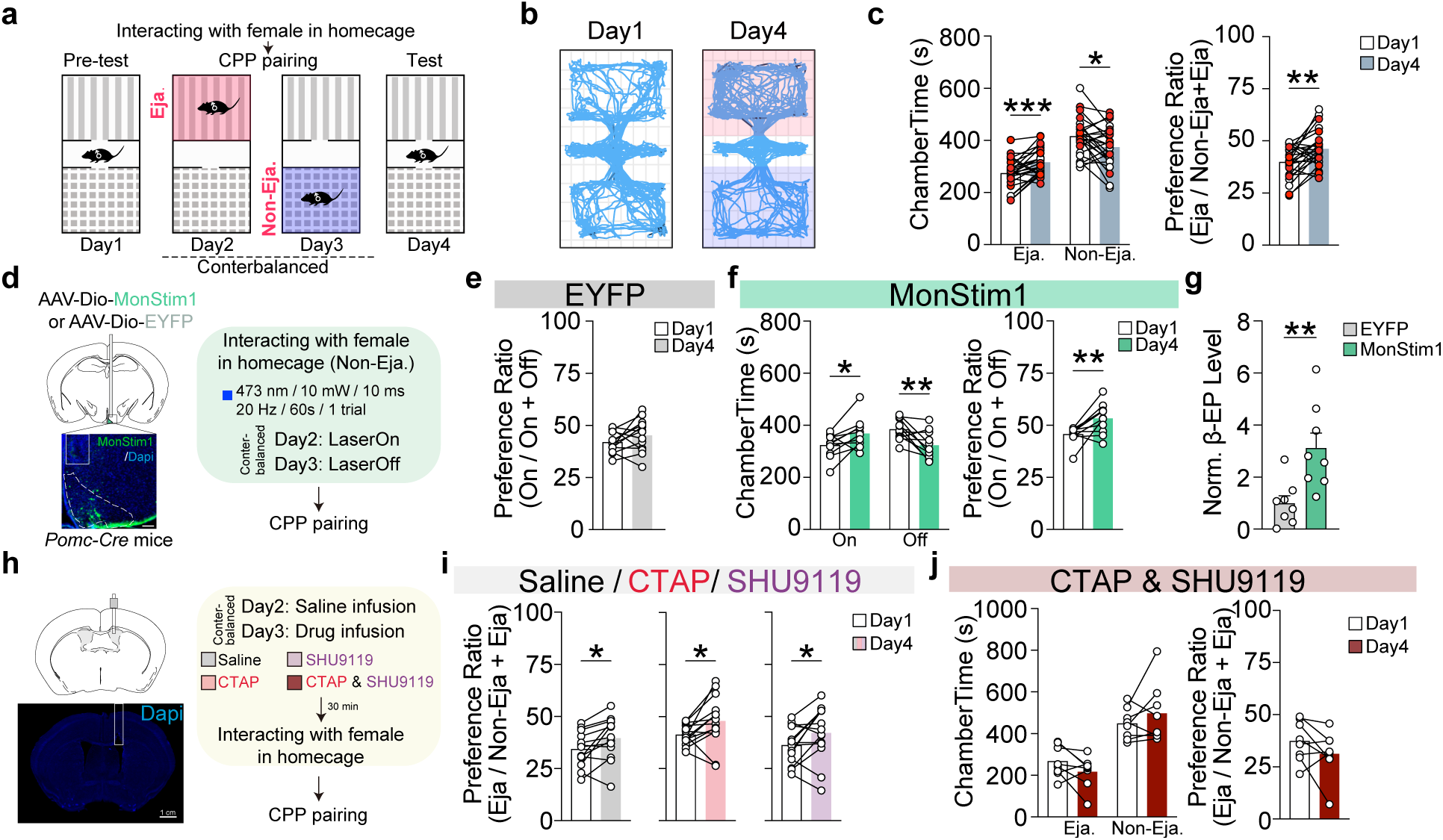
Regulation of ejaculation-conditioned place preference (CPP) by *Pomc+* neurons. **a.-b.** Ejaculation conditions CPP in wildtype male mice of both C57BL/6 and B6D2F1 hybrid background. **a.** the experimental procedure. Males were moved to one chamber achieving ejaculation (Eja) and the other chamber in not (Non-Eja); **b.** representative traces showing increased time spent in ejaculation-paired chamber after conditioning. **c.** Quantification of the time spent in each chamber (ChamberTime, left) and the preference score (right) for the ejaculation-paired chamber. Red circles and unfilled circles represent males of C57BL/6 and B6D2F1 hybrid background respectively. n = 12 C57BL/6 males and 13 B6D2F1 males. **d.** Schematics at the top (left) in shows AAVs encoding Cre-On MonStim1, or EYFP as control, were injected into the ARC of *Pomc-Cre* male mice of C57BL/6 background. The representative image at the bottom in shows ARC (outlined with dashed lines) MonStim1 expression, with solid lines indicating the implanted optic fiber. scale bar, 100 μm. Males interacted with a female in their home cages with the external fiber attached, without reaching ejaculation. On one of the CPP pairing days (day 2 or day 3), a single 60-second (473nm, 10mW, 10ms, 20Hz) laser stimulation was delivered. **e.-g.** Light delivery had no effects on the preference score for the light-paired chamber in the EYFP group (**e**), but significantly increased the time spent in, and the preference score for, the light paired chamber in the MonStim1 group (**f**). Additionally, light stimulation elevated CSF β-endorphin (β-EP) levels (**g**) in MonStim1 group. β-EP levels were normalized (norm.) to mean value of the EYFP group. n = 12 EYFP males for (e), 11 MonStim1 males for (f), 5 males/group for (g). **h.** Schematics at the top (left) and the presentative images at the bottom (left) show the implantation of a cannula in the lateral ventricle. Scale bar, 1cm. Saline, CTAP (μ-opioid receptor antagonist), SHU9119 (melanocortin receptor antagonist), or the combination of CTAP and SHU9119 were delivered through the implanted cannula into the lateral ventricle before the males interact with females in their home cages until ejaculation occurred prior to CPP pairing. Intromission pairing was preceded with saline perfusion. **i.-j.** Delivery of CTAP or SHU9119 alone could not block the ejaculation conditioned place preference (**i**) but the combination of CTAP and SHU9119 successfully blocked it (**j**). For each group, saline, n = 13; CTAP, n =14; SHU9119, n = 14; CTAP & SHU9119, n = 8. Values are presented as mean ± SEM. *p < 0.05, **p < 0.01, ***p < 0.001.

We next investigated whether artificial stimulation of *Pomc+* neurons could replicate the conditioning effects induced by ejaculation. To do this, we injected Cre-inducible AAVs encoding monStim1^38^, a light-sensitive Ca^2+^ modulator that elevates cytoplasmic Ca^2+^ levels via activation of Ca^2+^ release-activated Ca^2+^ (CRAC) channels, into *Pomc-Cre* males, followed by the implantation of an optical fiber (**Fig. 5d**). As the control, AAVs encoding Cre-inducible EYFP were injected into *Pomc-Cre* males (**Fig. 5d**). In these experiments, males were allowed to interact with female in their home cages without reaching ejaculation, with an optical fiber attached (**Fig. 5d**). On one day, a single light stimulation (473 nm, 10 mW, 10 ms, 20 Hz, 60 s) was delivered, while on the other day, no light was applied. Afterward, the males were immediately transferred to one of two chambers of the CPP apparatus for pairing (**Fig. 5d**).

As expected, light stimulation in the EYFP control group had no effect on the time spent in, or the preference score for, the light paired chamber (**Fig. 5e**). In contrast, optogenetic stimulation of *Pomc+* neurons in the monStim1 group increased the time spent in the light-paired chamber, resulting in a higher preference score (**Fig. 5f**), thereby mimicking the rewarding effects of ejaculation. *Post-hoc* analysis further revealed that CSF β-endorphin levels were elevated in the mon Stim1 group after light stimulation (**Fig. 5g**). A previous study demonstrated that chemogenetic activation of *Pomc+* neurons is mildly aversive^39^. In contrast, our findings show that optogenetic activation of *Pomc+* neurons in the context of male mating not only increases β-endorphin levels but also replicates the place-conditioning effects associated with ejaculation.

Conversely, we explored whether blocking the signaling of neuropeptides secreted by *Pomc+* neurons could prevent the place preference conditioned by ejaculation. To achieve this, we implanted males with a cannula for drug infusion into the lateral ventricles and repeated the CPP protocol (**Fig. 5h**). In these experiments, males were injected intracerebroventricularly (i.c.v.) with saline or drugs that block neuropeptide receptor signaling, including SHU9119 (a melanocortin receptor antagonist)^40^ and CTAP (a selective, competitive μ-opioid receptor antagonist)^41^, and then allowed to mate with a female in their home cages until ejaculation (**Fig. 5h)**. Alternatively, they were injected i.c.v. with saline and allowed to interact with female for a similar duration without reaching ejaculation. Immediately after mating, the males were transferred to one of two chambers for pairing, as previously described (**Fig. 5h**).

We found that infusion of saline or individual receptor blockers (SHU9119 or CTAP) did not affect the preference score for the chamber associated with ejaculation (**Fig. 5i**). However, simultaneous infusion of both CTAP and SHU9119, which blocks both μ-opioid receptor and melanocortin receptor signaling, successfully prevented the development of place preference for the ejaculation-paired chamber (**Fig. 5j**).

These findings collectively indicate that the *Pomc* neuronal activity are sufficient whereas signaling of *Pomc* neuropeptides are necessary for the sexual reward associated with ejaculation. In summary, we have identified an intra-hypothalamic circuit from POA *Calb1+* neurons to arcuate *Pomc+* neurons that coordinates the release of β-endorphin with ejaculation, illustrating the neurobiological mechanisms underlying the post-ejaculation emotional state.

## Discussion

Although circumstantial evidence has implicated the endogenous opioid system in encoding sexual reward^2,3,13,18,19,36,42–44^, the specific role of *Pomc+* neurons—an important source of endorphins in the brain—in male mating behavior remained largely unexplored, especially compared to the well-established roles of these neurons in feeding and metabolism^33,45^. In this study, beginning with the identification of ejaculation-specific activation of POA *Calb1+* neurons, we identified a distinct pattern of sustained Ca^2+^ transients in *Pomc+* neurons downstream of the POA specifically during ejaculation, which corresponds with increased β-endorphin levels in male mice post-ejaculation. Additionally, we showed that artificially activating *Pomc+* neurons increased CSF β-endorphin levels and conditioned place preference similar to that induced by ejaculation. Conversely, blocking *Pomc* neuropeptide signaling abolished the place conditioning effects induced by ejaculation. These findings collectively establish a distinct role for *Pomc+* neurons in encoding the affective state following ejaculation.

The ejaculation-associated activity in *Pomc+* neurons persisted for an average of 50 seconds, significantly exceeding both the motor action of ejaculation itself and the activity observed in POA *Calb1+* neurons. Notably, *Pomc+* neurons did not exhibit any activity when ejaculation was induced via electrical pulses delivered through a rectal probe (Extended Data Fig. 9a-9c). This indicates that *Pomc+* neuron activation is not solely responsive to the motor aspects of ejaculation but requires a specific behavioral context. Along this line, we suspect that stimuli such as light touch or mechanic vibration applied to the penis, or optogenetic activation of Krause corpuscle afferent terminals alone – while known to induce penile erection^32^ necessary for vaginal penetration and intromission – are unlikely to trigger the sustained *Pomc+* neuron activation observed during ejaculation.

As such, the prolonged activity of *Pomc+* neurons during ejaculation suggest a role in encoding a distinct internal affective state. Interestingly, previous studies have shown that transient exposure to predator cues elicits prolonged activation of *SF1+/Nr5a1+* neurons in the dorsomedial and central subdivisions of the ventromedial hypothalamus (VMHdm/c)^46^, with a duration comparable to the ejaculation-related activity in *Pomc+* neurons. Neural recordings at single-cell level reveal that the sustained firing of VMHdm/c *SF1+/Nr5a1+* neurons at the population level emerge from individual neurons that responded with distinct temporal dynamics^46^. Theoretical models indicate that both recurrent local excitatory synaptic connections and slow neuronal dynamics, mediated by neuropeptide signaling, are necessary to produce persistent activity at the population level^46^. Further research indicates that the persistent firing of VMHdm/c *SF1+/Nr5a1+* neurons encodes a defensive internal state rather than specific defensive behaviors^46,47^.

Given the similarities in the duration of activity and their proposed roles in encoding emotional states, it is plausible that similar cellular mechanisms may also underlie the persistent ejaculation-associated activity observed in *Pomc+* neurons. While recurrent connections and local neuropeptide signaling among *Pomc+* neurons have been suggested^48^, experimental evidence remains inconclusive. Exploring whether and how these local signaling mechanisms contribute to the sustained firing observed in *Pomc+* neurons during ejaculation is an exciting avenue for future research. Additionally, considering the prevalence of recurrent connections and neuropeptide expression in the hypothalamus^49,50^, persistent firing of distinct hypothalamic neurons may play a generalized role in encoding different emotional states^51–54^.

Regardless of the local mechanisms, we demonstrate that POA *Calb1+* neurons, which are upstream of *Pomc+* neurons and exhibit ejaculation-specific activity, likely serves as a trigger for sustained *Pomc+* neuron activity and β-endorphin release during ejaculation. This conclusion is supported by evidence showing that the ablation of POA *Calb1+* neurons results in a significant reduction, approximately 30%, in the number of c-Fos*+* signals detected in *Pomc+* neurons following ejaculation. Interestingly, while the inhibition or ablation of POA *Calb1+* neurons delays ejaculation and extends the mating process, it does not entirely prevent ejaculation. This partial effect may result from insufficient inhibition of POA *Calb1+* neurons. Alternatively, other neural populations might contribute to the transition from intromission to ejaculation in the absence of POA *Calb1+* neurons. Specifically, other neurons providing inputs to *Pomc+* neurons (Extended Data Fig. 9d-9g) could also play a role in regulating ejaculation timing, ensuring coordination of ejaculation with *Pomc+* neuron activation, even without inputs from POA *Calb1+* neurons. This hypothesis merits further investigation.

Finally, *Pomc+* neurons are believed to secrete various neuropeptides, including β-endorphin^5^. While this study did not specifically examine other neuropeptides, it is plausible that additional neuropeptides are also released by *Pomc+* neurons during sustained activation following ejaculation. These neuropeptides could enter the cerebrospinal fluid (CSF) or be released post-synaptically, potentially playing important roles in mediating post-ejaculatory affective states. This is further supported by findings that blocking the μ-opioid receptor alone is insufficient to eliminate the place-conditioning effects associated with ejaculation; simultaneous blockade of Melanocortin receptors is also required. Given the evolutionary conservation of the hypothalamus, hypothalamic neuropeptides, and male mating behaviors across mammals, we anticipate that similar mechanisms may mediate post-ejaculatory affective states in non-rodent species. Supporting this idea, there is circumstantial evidence suggesting that opioid and melanocortin pathways are involved in male sexual behaviors and post-ejaculatory affective states in humans^2,3,13^.

In conclusion, we have identified a distinctive activity pattern of *Pomc+* neurons coinciding with ejaculation. These findings enhance our understanding of *Pomc+* neuron function and pave the way for future studies into post-ejaculatory affective states and related disorders.

## Materials and Methods

### Animals

C57BL/6 wild-type animals were obtained from SLAC Laboratory Animal (Shanghai). B6D2F1 wild-type males were purchased from Beijing Vital River Laboratory Animal Technology Co., Ltd. Genetically modified animals, including *Calb1-Cre* (Calb1^tm2.1(cre)Hze/J^, #028532), *Pomc-Cre* (Tg(Pomc1-cre)16Lowl/J, #005965), *Agrp-Cre* (Agrp^tm1^(cre)^Lowl/J^, #012899), and *Adcyap1-Cre* (Adcyap1^tm1.1(cre)Hze/ZakJ^, #030155), were purchased from the Jackson Laboratory. These animals were housed at the Center for Excellence in Brain Science and Intelligence Technology (CEBSIT) animal facility. All Cre lines were backcrossed onto the C57BL/6 background for more than two generations unless stated otherwise. To produce *Pomc-Cre* animals with a hybrid background, *Pomc-Cre* females on the C57BL/6 background were mated with DBA mice, purchased from Beijing Vital River Laboratory Animal Technology Co., Ltd. Only heterozygous animals for Cre alleles were used in the study. Only adult animals (8-24 weeks for males, and 6-24 weeks for females) were used. Animals were maintained on a reversed 12 h:12 h light/dark cycle with food and water provided *ad libitum*. Littermates were randomly assigned to the experimental or control group. All experimental protocols were approved by the Animal Care and Use Committee of CEBSIT, Chinese Academy of Sciences, Shanghai, China (IACUC No. NA-016-2016).

### Virus

The following viruses were used in this study: AAV-hSyn-DIO-TVA-mCherry-2A-oG (serotype 2/9, titer 4.8×10^13^ vg/mL), AAV-hSyn-DIO-EGFP-cry2-Mon-Stim1 (serotype 2/8, titer 1.08×10^13^ vg/mL) were purchased from CEBSIT gene editing core facility. AAV-EF1a-DIO-RVG (serotype 2/9, titer 2.0 ×1012 vg/mL), AAV-EF1a-DIO-EGFP-2A-TVA (serotype 2/9, titer 2.0 ×1012 vg/mL), RV-EnVA-deltaG-dsRed (2× 10^8^ infectious units / ml) were purchased from BrainVTA, Wuhan. AAV-hSyn-DIO-GCamp6s (serotype 2/8, titer 8.0×10^12^ vg/mL), AAV-hSyn-DIO-EYFP (serotype 2/8, titer 5×10^12^ vg/mL), AAV-hSyn-DIO-mCherry (serotype 2/5, titer 3.5×10^12^ vg/mL), AAV-hSyn-FAS-GCamp6s (serotype 2/9, titer 7.0×10^12^ vg/mL), AAV-hSyn-FAS-EYFP (serotype 2/9, titer 4.0×10^12^ vg/mL), AAV-hSyn-DIO-ChR2-mCherry (serotype 2/9, titer 6.0×10^12^ vg/mL), AAV-hSyn-FAS-ChR2-mCherry (serotype 2/9, titer 6.0×10^12^ vg/mL), AAV-hSyn-DIO-NpHR3.0-mCherry (serotype 2/9, titer 3 ×10^12^ vg/mL), and AAV-CAG-DIO-taCaspase3-TEVp (serotype 2/5, titer 8.5 ×10^12^ vg/mL) were purchased from Taitool Bioscience, Co., Shanghai. Rabies-SADδG-GFP-EnVA virus was generated as described before^55,56^. Briefly, rabies-SADδG -GFP viruses (RVdG-GFP) were amplified from local viral stocks in B7GG cells (kindly provided by Dr. Ed Callaway, Salk Institute). EnVA pseudotyped rabies was generated in BHK-EnVA cells. The virus was concentrated by two rounds of centrifugation, suspended in Hank’s Balanced Salt Solution (GIBCO), and titered in HEK293-TVA cells (kindly provided by Dr. J. A. T. Young, Salk Institute) with serial dilutions of the virus. The titers of the RVdG-GFP-EnVA were in the range of 10^7^ – 10^8^ infectious units / mL. The virus was stored at -80℃ until use.

### Stereotactic surgery

All surgeries were performed as previously described^8^. Animals were anesthetized under isoflurane (0.8% - 5%) and mounted to a stereotaxic frame (David Kopf Instrument, Model 1900). Surgeries were performed with steady isoflurane (1-2%) in mixed air (95% O_2_, 5% CO_2_). For viral injection, the skull was exposed with a small incision, and holes were drilled for virus delivery with glass pipettes (15 - 25μm in diameter at the tip). The following coordinates, according to the standard mouse brain atlas (Paxinos and Franklin Mouse Brain Atlas, 2nd edition), were used to target the arcuate nucleus, AP: - 1.85 mm, ML: ± 0.25 mm, DV: - 5.85 mm; POA, AP: + 0.10 mm, ML: ± 0.40 mm, DV: - 5.15 mm. We injected 200 - 400 nL virus into each site at 80-100 nL / min with a homemade injector (adapted from MO-10 Narishige, Japan). The pipette was left in place for about 10 mins after the injection for virus diffusion.

For fiber photometry recording experiments, the virus was unilaterally injected, and an optic fiber (diameter, 200 μm; N.A., 0.37; AniLab Software and Instruments Co., Ltd) was implanted 50 μm above the coordinates of the virus injection site. For optogenetic activation, the virus was unilaterally injected, and an optic fiber (diameter, 200 μm; N.A., 0.37; AniLab Software and Instruments Co., Ltd) was implanted 200 μm (ARC), or 400 μm (POA) above the coordinates of the virus injection site. For optogenetic inhibition, the virus was bilaterally injected, and dual optic fibers (DFC_200/245–0.37_6.0mm_DF0.9_FLT; Doric Lense) were implanted 400 μm (POA) above the coordinates of the virus injection site. The implanted optic fibers were secured onto the skull with adhesive MetaBond (C&B MetaBond).

For pharmacological experiments, a cannula was implanted for drug infusion in the lateral ventricle. A single guide cannula (#62004, RWD Life Science) was unilaterally implanted to target either the left or right lateral ventricle with the coordinate AP: - 0.30 mm, ML: ± 1.00 mm, DV: - 1.50 mm. The guide cannula was inserted with a dummy cannula (#62108, #62168, RWD Life Science) to prevent clogging during the recovery period and secured with a dust cap.

For neural tracing experiments (Fig. 3a-3b, Extended Data Fig. 8), the virus was bilaterally injected. For pseudorabies tracing experiments, approximately 150 nL of a 1:1 mixture of helper viruses (AAV-EF1a-DIO-RVG and AAV-EF1a-DIO-EGFP-2A-TVA) or 150 nL of AAV-hSyn-DIO-TVA-mCherry-2A-oG alone was unilaterally injected into the ARC of *Pomc-Cre* mice. Three weeks later, ∼100–150 nL of RV-EnVA-deltaG-DsRed or Rabies-SADδG-GFP-EnVA was injected into the same location.

### Female receptivity induction

C57BL/6J wild-type female mice were surgically ovariectomized as adults (6-8 weeks age) and hormonally primed to induce receptivity. Hormones were suspended in sterile sunflower seed oil (Sigma-Aldrich, S5007). We injected 10 μg (in 50 μL oil) and 5 μg (in 50 μL oil) of 17β-estradiol benzoate (Sigma-Aldrich, E8875) 48 h and 24 h preceding the mating test. On the test day, females were injected with 50 μg of progesterone (Sigma-Aldrich, P0130; in 50 μL oil) 4 – 6 h prior to the mating test. All the females were hormone primed and trained at least twice with males to obtain sexual experience before being used in experiments. Females were given at least a seven-day interval between two mating test trials.

### Male mating behavior test

All males, except those used for optogenetic activation experiments in Extended Data Fig. 5, had prior sexual experience with females. This experience consisted of two sessions of either overnight co-housing with a female or training during a mating behavior test where the males achieved ejaculation. Details regarding the males’ sexual experience are provided in Extended Data Table 1. Males were single-housed for 5–7 days before the behavior test. The tests began at least one hour after the onset of the dark cycle and were recorded using an infrared camera at a frame rate of 25 Hz. Mice were first transferred to the test room in their home cages and allowed to acclimate for at least 10 minutes. For each mating trial, a hormone-primed female was introduced into the male’s home cage, where they interacted freely for 30-60 minutes, unless otherwise specified. Afterward, the female was removed. Mating tests were conducted 1– 4 times for each male, with a minimum interval of three days between trials, except for those conducted before the CPP experiments.

Videos were manually annotated frame-by-frame using a custom MATLAB program, as previously described^8^. Specific male-initiated actions, including genital sniff, mount, intromission, and ejaculation, were defined and annotated by experimenters blinded to the experimental group information. “Genital sniff” was defined as an anogenital investigation when the male initiated nose-to-anogenital contact. “Mount” was scored when the male placed its forelimbs on the female’s back, climbed on top, and moved its pelvis. Rhythmic pelvic movements following a mount were scored as “Intromission”. “Ejaculation” was scored when the male displayed a deep thrust followed by firmly seizing the female in a stereotyped, immobile posture after intromission, as shown in Extended Data Video 1-4. The onset of mount is defined as the first frame when the male grasped and mounted the female’s flanks by his forelimbs; the onset of intromission is defined as the first frame when the male began rapid and rhythmic pelvic thrust; the onset of ejaculation is defined as the first frame of the last deep thrust.

### Single-nucleus RNA sequencing

#### Sample preparation, library generation, and sequencing

For sample preparation for the Act-Seq (Fig. 1), males were tested in parallel in the same room, following a similar handling procedure. A hormone-primed female mouse was introduced into the males’ home cages. The behavioral test was terminated once one of the tested males achieved ejaculation or more than five intromissions. The animals that ejaculated were assigned to the “ejaculation” group, while those achieved intromission but not ejaculation were assigned to the “intromission” group. All animals were sacrificed approximately 30 minutes after the behavioral test ended.

Mice were anesthetized with isoflurane, perfused trans-cardinally with ice-cold oxygenated (95% O_2_/5% CO_2_) cutting solution (in mM: 2.5 KCl,1.25 NaH_2_PO_4_, 2 Na_2_HPO_4_, 2 MgSO_4_, 213 sucrose, 26 NaHCO_3_). The brain was promptly placed in ice-cold oxygenated cutting solution and sectioned at 250 μm using a vibratome (VT1200S, Leica). Immediately after the sectioning, brain slices were transferred to the dissection chamber with ice-cold oxygenated ACSF (in mM: 126 NaCl, 2.5 KCl, 1.25 NaH_2_PO_4_, 1.25 Na_2_HPO_4_, 2 MgSO_4_, 10 glucose, 26NaHCO_3_, 2 CaCl_2_) containing Actinomycin D (1 μM, CST, 15021S). Then, the preoptic area (POA) was carefully dissected according to the Allen brain atlas (www.brain-map.org) and collected to the corresponding tubes that were stored on the dry-ice. After the collection of POA from all qualified mice for either group, tubes were put into the liquid nitrogen before being processed further for library construction and sequencing. The durations of the dissection procedure for each animal were less than 10 min from the time point of perfusion to frozen in the tube. Three mice were included in either group. Subsequent steps, including the preparation of single-nucleus suspensions, library construction, and sequencing, were carried out by BGI Genomics using the Chromium Next GEM Single Cell 3’ Kit V3.1 (10X Genomics) in accordance with the manufacturer’s protocol.

#### Normalization, clustering, and differential gene expression

Raw sequencing files were aligned to the mm10 mouse genome (GRCm38) and converted to gene expression matrices using the Cell Ranger pipeline (Cell Ranger 7.2.0) with default parameters. Intronic reads were included to enhance assay sensitivity.

All downstream analyses of scRNA-seq data were performed using the R package Seurat (v4). First, the scRNA-seq datasets were merged to generate the integrated file, and the quality of the sequenced cells was assessed. Outlier cells were identified based on the following criteria: 1) fewer than 200 detected genes (indicating potential fragment or dead cells); 2) the UMI count of mitochondrial genes exceeding 5% (indicating potential dead cells); 3) more than 6,000 genes (indicative of multiplets), and were filtered out. To focus on neurons, coarse preliminary clustering was performed to define major cell types. Clusters that lack expression of neuronal genes including *Snap25, Stmn2, Gad1/Gad2/Slc17a6* or those that express conventional non-neuronal cell markers including *Gfap, Pdgfra, Aqp4, C1qc, Gja1, Cldn5, Opalin, Mustn1* were excluded from further analysis. The final dataset comprised 27,447 neuronal nuclei, with 15,453 from the intromission group and 11,768 from the ejaculation group.

The expression of each gene in each cell was normalized by total number of UMI counts detected in that cell, then multiplied by a scale factor of 10,000 prior to log transformation (NormalizeData function). Gene expression values were then scaled by mean/variance across cells to range between 0 and 1, and the proportion of mitochondrial UMIs were regressed out (ScaleData function). Clustering using FindClusters function yielded unique transcriptomic clusters. The analysis and clustering procedures were as follows: Firstly, highly variable genes were identified (FindVariableGenes function; top 3,000 genes with the highest standardized variance selected by selection.method =‘vst’) and used as input for dimensionality reduction via principal component analysis (PCA) after removing immediate early genes (e.g., *Fos*, *Fosl2*, *Junb*, *Egr1*, *Arc*, *Homer1*; 139 genes in total as previously described^25^). Secondly, the PCs from previous step were used for clustering analysis (FindClusters function). Thirdly, differentially expressed genes in subclusters were detected using the FindAllmarkers function.

To identify neurons activated under ejaculation or intromission conditions, neurons with the IEGs’ count greater than 0 were classified as IEGs+, while all other neurons were classified as IEGs–. To identify activated neuronal clusters, the enrichment score of IEG+ neurons relative to IEG-neurons for each immediate early genes (IEGs) within each neuronal clusters was calculated using a binomial test for the ejaculation or intromission conditions separately, with false discovery rate (FDR) correction, and plotted in Fig. 1d and Extended Data Fig. 2.

### Fiber photometry recording

Fiber photometry recordings were conducted as previously described^8^. Briefly, the virus was allowed to express for 3 to 4 weeks before recording. Males were singly housed for 3 to 7 days before fiber photometry recording and were trained by co-housing with hormone-primed females overnight twice to gain sexual experience. Before each recording session, the implanted optic fiber was connected to the recording setup (Biolink Optics Technology Inc., Beijing) via an external patch cord. A 488 nm excitation laser (OBIS 488LS; Coherent) was reflected off a dichroic mirror (MD498, Thorlabs) and directed to the neural tissues through the implanted optic fibers. After the patch cord was attached to the optic fiber implanted in the mice, they were allowed to move freely and acclimate to the device for 5 minutes.

During signal recording, males were first recorded for 3 minutes to collect fluorescent baseline signals. The detected fluorescence was filtered with a band pass filter (MF525-39, Thorlabs) and collected by a photomultiplier tube (PMT, R3896, Hamamatsu). Emission signals were low-pass filtered at 30 Hz and sampled at 500 Hz using a data acquisition card (USB6009, National Instrument) with software provided by Biolink Optics. Subsequently, a hormonally primed female was introduced into the male’s cage, and fluorescent signals were recorded until the male achieved ejaculation or 60 minutes had elapsed since the female’s introduction.

To analyze signals from fiber photometry experiments, raw fluorescence signals were adjusted for overall trends to account for photobleaching. Animals showing jitter, square wave signals, or low signal-to-noise ratios were excluded from further analysis. Behavior annotations were consistent with those used in the mating test. To align fluorescence signals with behaviors, data were segmented based on behavioral events within individual trials, and average signals of 10 to 15 seconds before behavior initiation were used as F0. Fluorescence changes (ΔF/F) were calculated as (F−F0)/F0 and averaged first across trials and then across animals. For statistic comparison, the average (ΔF/F) signals during behavior were compared to 0 using a one-sample t-test to determine statistical significance.

### Optogenetic Inhibition

A bilateral bundled optic fiber (Inper, Co., Ltd) was used to connect a 589 nm laser power source (Changchun New Industries Optoelectronics Tech Co., Ltd.) to the implanted optic fiber in the animal. The external optical fiber was attached to a rotary joint (Doric Lense) to allow the animal to move freely with attached fibers. The tested animal was allowed to first explore the cage for about 10 to 15 min with the external fiber attached. After that, a hormonally primed female was introduced. Then, a custom-written program in MATLAB was started to control a Master 9 (A.M.P.I.), which sent a trigger signal to initiate recordings from the camera. Yellow light (∼ 6 mW) was automatically triggered and continuously delivered whenever the tested animal was detected within one body length (∼ 9 cm) of the female during the 30-minute mating behavior tests. Light inhibition trials and no-light trials were alternated randomly for each animal.

### Optogenetic Activation

An optic fiber (Inper, Co., Ltd) was used to connect a 473 nm laser power source (Shanghai Laser and Optics Century Co.) to the implanted optic fiber in the animal. The external optical fiber was attached to a rotary joint (Doric Lense) to allow the animal to move freely with the fibers attached. The animal was first allowed to explore the cage for about 10 to 15 minutes with the external fiber attached. Afterward, a hormonally primed female was introduced. A custom-written MATLAB program was then initiated to control a Master 9 (A.M.P.I.), which sent a trigger signal to start camera recordings and controlled the laser to deliver eight 15 seconds of photo-stimulation at 20Hz, 12 mW, spaced randomly 90–120 seconds apart (POA activation), or a single 60-second photo-stimulation at 20Hz, 12 mW (ARC activation).

### Conditioned Place Preference (CPP)

The CPP apparatus is an acrylic box consisting of three compartments, including a neutral middle zone (75 x 190 x 145 mm) and two distinct conditioning chambers (190 x 190 x 145 mm), with one featuring a mesh wire floor and white walls, and the other a slatted wire floor and black walls, providing distinct visual and tactile cues. The CPP apparatus was placed in a video-recording box illuminated by dim white light. Males interacted with a hormonally primed female in their home cage, within a similar video-recording box without light, before being placed into the CPP apparatus for conditioning.

The CPP procedure consisted of three phases over 4 days: pre-test, conditioning, and test. On day 1, during the pre-test, mice were introduced into the middle chamber and allowed to explore all three chambers freely for 15 minutes. The time spent in each chamber was recorded to identify the non-preferred and preferred chambers. On days 2 and 3, during the conditioning phase, mice underwent mating behavior testing and were immediately confined to one of the conditioning chambers for 30 minutes in a counterbalanced manner. On day 4, during the test phase, mice were placed in the middle zone and allowed to freely explore all chambers for 15 minutes. The time spent in each paired chamber was manually recorded and used to calculate a preference ratio, determined by dividing the time spent in the paired chamber by the total time spent in both conditioning chambers.

For ejaculation conditioning, mice reached ejaculation on one of the conditioning days and were confined to the initially less preferred chamber. On the other day, when they did not reach ejaculation, they were confined to the preferred chamber. For conditioning with optogenetic stimulation of *Pomc+* neurons, animals, connected with an optic fiber, and a single trial of blue laser (10 mW, 20 Hz, 10 ms, 60 s) was delivered. On this day, mice were confined to the initially less preferred chamber, while the other day when no light was delivered the mice were confined to the preferred chamber.

For the pharmacological inhibition test, the CPP procedures were the same as the ejaculation-conditioning protocol, with the addition of saline or drug injections before the mating test on the conditioning day. Drugs, including SHU9119 (Tocris, Cat#3420), CTAP (Tocris, Cat#1560), or a mixture of the two, were administered prior to ejaculation pairing. Briefly, a dummy injector was inserted into the guide cannula to ensure a clear passage, and then removed. Drugs were microinjected into the lateral ventricle using a single injector (#62204, RWD Life Science) for intracerebroventricular (i.c.v.) injection. The injector extended 1 mm beyond the tip of the guide cannula and was connected by PE tubing to a 1 or 2 μL microinjection syringe mounted on a pump (Harvard Apparatus). Drugs included SHU9119 (0.5 nmol), CTAP (0.3 nmol), a mixture of both in 2 μL volume, or 2 μL of saline as a control. They were all injected at a rate of 30 nL/s. After injection, the injector was left in place for 30 seconds to allow drug diffusion before removal. The dummy injector and cap were then replaced, and the mice were returned to their home cages to undergo the mating test and conditioning.

### CSF collection and endorphin measurement

Cerebrospinal fluid (CSF) was collected from mice following the protocol described previously^57^. A sharpened glass capillary with a tip diameter of approximately 10-20 µm was used for CSF collection. The capillary was connected to a syringe via a thin tube and a three-way valve, which allowed positive pressure to be applied to expel any contaminants. Mice, including those in the post-ejaculation, post-intromission, and opto-activation groups, were immediately anesthetized with 4% chloral hydrate. Afterward, the hair was removed from the back of the mouse’s head and the mouse’s head was positioned at a 45° angle, an incision was made to expose the base of the skull. The dura mater over the cisterna magna was then cleaned and dried, allowing approximately 10 uL of CSF to be collected using the prepared capillary. For some mice, after CSF collection, the wound was sutured, and the animals were allowed to recover for a week before a second collection. Each animal underwent no more than two CSF collections. CSF samples were stored at 4°C and processed with the ELISA test within a week or were stored at -20°C until testing. Endorphin levels were measured using the Mouse Beta-Endorphin, B-EP ELISA Kit (CUSABIO, Cat# CSB-E06827m), following the manufacturer’s protocols. Relative endorphin levels were normalized to the mean endorphin level of animals in either the intromission or the *Pomc-Cre*-EYFP group within each batch.

### Immunohistochemistry

Animals were anesthetized with 0.2 mL 10% chloral hydrate in sterile saline and perfused with 15 mL PBS, followed by 15 mL ice-cold 4% PFA. Brains were dissected and post-fixed overnight in 4% PFA at 4℃. Brains were sectioned at 40 μm using a vibratome (VT1000S, Leica). Sections were washed with PBS and blocked with blocking reagent (5% goat serum, 0.1% Triton X-100, 2 mM MgCl_2_ in PBS) for 1 hour at room temperature. Then sections were incubated in AGT solutions (0.5% goat serum, 0.1% Triton X-100, 2 mM MgCl_2_ in PBS) with appropriate primary antibody overnight at 4℃. On the second day, slices were washed with AGT solutions three times and incubated in AGT solutions with appropriate secondary antibodies for 2 hours at room temperature. Slices were then rinsed in AGT solutions and counterstained with DAPI (dilution 1:1,000, Sigma, 5 mg / mL, Cat #d9542) for 10 min. Then, slices were washed in PBS solutions and mounted to slides. For GCaMP6s and EYFP staining, the primary antibody was anti-GFP antibody (dilution 1:1,000, ABCAM, Cat #ab13970), and the second antibody was goat-anti-chicken conjugated with Alexa fluor 488 (Jackson ImmunoResearch, Cat# 103-545-155, 1:1,000). For mCherry staining, the primary antibody was anti-dsRed antibody (dilution 1:1000, Clontech, Cat #632496), and the second antibody was goat anti-rabbit conjugated with Cy3 (Jackson ImmunoResearch, Cat# 111-165-003, 1:1,000). For Calb1 staining, the primary antibody was anti-Calbindin1 antibody (dilution 1:1,000, swant, Cat #CB-38a), and the second antibody was goat anti-rabbit conjugated with Cy3 (Jackson ImmunoResearch, Cat# 111-165-003, 1:1,000). For ACTH staining, the primary antibody was anti-ACTH/CLIP (dilution 1:100, santa cruz, Cat #373878), and the second antibody was goat anti-mouse conjugated with 647 (Jackson ImmunoResearch, Cat# 115-605-003). For c-Fos staining, the primary antibody was anti-c-Fos (dilution 1:1000, synaptic system, Cat #226308, 1:1,000), and the second antibody was goat anti-Guinea pig conjugated with Cy3 (Jackson ImmunoResearch, Cat# 111-165-003, 1:1,000). Images were captured by a 10 X objective on a fluorescent microscope (Olympus, VS120) or a confocal microscope (Nikon, C2). Images were processed and counted using the NIH ImageJ software. All analyses were done by an experimenter blind to the genotype and the drug treatment information.

### In situ hybridization

DNA templates for generating in situ *Calb1* probes were cloned using the following primer sets: Forward 5’-gaactattcaggatgtgtggca -3’ and Reverse 5’-gggctatggtcatactctctgg-3’. Anti-sense RNA probes were transcribed with T7 RNA polymerase (Promega, Cat# P207E) and digoxigenin (DIG)-labeled nucleotides. Animals were anesthetized with 10% chloral hydrate and transcardially perfused with DEPC-treated PBS (DEPC-PBS) followed by ice-cold 4% paraformaldehyde (PFA). Afterward, brains were sectioned at 40 μm thickness using a vibratome (VT1000S, Leica). Brain sections were washed in 2XSSC buffer containing 0.1% triton for 30 min, acetylated in 0.1 M triethanolamine (pH 8.0) with 0.25% acetic anhydride (vol/vol) for 10 min, equilibrated in pre-hybridization solution for 2 h at 65 ℃ and subsequently incubated with 0.5 μg/ mL of *Calb1* probes in hybridization buffer overnight at 65 ℃. The next day, sections were rinsed in pre-hybridization solution and pre-hyb/TBST (TBS with 0.1% tween-20) for 30 min each. Next, sections were washed with TBST for twice and TAE for three times, each for 5 min. Sections were then transferred into wells in 2% agarose gel, which were run in 1XTAE at 60 V for 2 h to remove unhybridized probes. Sections were then washed twice in TBST, and subsequently incubated with sheep anti-digoxygenin-AP (1:2000, Roche, Cat# 11093274910) in 0.5% blocking reagent (Roche, 11096176001) at 4 °C overnight. On the second day, sections were washed and stained with NBT (Roche, Cat# 11383213001) and BCIP (Roche, Cat# 11383221001) for 3 h at 37 ℃. All sections were washed after staining and mounted on glass slides. Images were captured with ×10 objective using a conventional microscope.

### RNAscope

Animals were anesthetized with 0.2 mL of 10% chloral hydrate in sterile saline and perfused with 15 mL of DEPC-PBS, followed by 15 mL of ice-cold 4% PFA. Brains were dissected and post-fixed overnight in 4% PFA at 4 °C, then dehydrated in 30% sucrose in DEPC-PBS overnight. Subsequently, brains were sectioned at a thickness of 20 μm and mounted onto SuperFrost Plus® Slides (Fisher Scientific, Cat. No. 12-550-15). After air-drying, slides were stored at −80 °C until processing according to the RNAscope® Multiplex Fluorescent Reagent Kit v2 User Manual (ACD Bio.). Probes for *Calb1* (Lot#428431-C2), *Adcyap1* (Lot#409511-C1), *Fos* (Lot#316921-C3), and *Pomc* (Lot#314081-C1) were ordered from ACD Bio. and used in the experiment.

For RNAscope analysis combined with immunohistochemistry, after the completion of RNAscope procedures, brain slices were blocked for 1 hour at room temperature in PBS with 2.5% BSA, followed by incubation with the primary antibody chicken anti-GFP (1:300, Abcam, Ab13970) at 4 °C overnight. The secondary antibody goat anti-chicken IgG-Alexa 488 (1:300, Jackson ImmunoResearch Laboratories, 103-545-155) was applied in PBS with 1.25% BSA at room temperature for 2 hours. Brain sections were then rinsed three times with PBS and counterstained with DAPI (Sigma, Cat# d9542, 5mg/mL, dilution 1:1000). All images were captured using a 20× objective on a confocal microscope and processed and counted using NIH ImageJ software.

### Quantification of axonal projection and RV inputs

To quantify the axonal projection, every other section of coronal serial sections of 40 μm thickness was imaged using a conventional microscope with a 10x objective lens (0.6499 mm / pixel, VS120, Olympus). To correct for the non-specific background noise of each brain section, the whole image of the brain sections were first evened using a custom written MATLAB (MathWorks) code that extract pixels with value between 2 standard deviation of the mean intensity. The average pixel intensity in a target brain region containing ChR2-mCherry terminals signals was then corrected by subtracting the background intensity value obtained from an adjacent brain region that did not contain mCherry+ signals. This corrected value was then normalized to the pixel intensity of the POA containing the ChR2-mCherry+ neuronal soma for each animal.

To quantify the number of retrogradely labeled RV-dsRed neurons, every other coronal section (40 μm thickness) was imaged using a conventional microscope (10x objective lens, 0.6499 mm/pixel, VS120, Olympus). Labeled cells in each region of interest were manually counted by experimenters. Brain areas were identified using DAPI immunostaining and the mouse brain atlas (Paxinos and Franklin’s The Mouse Brain in Stereotaxic Coordinates). The number of RV labeled neurons in upstream brain regions were normalized to the number of starter neurons (GFP+/dsRed+ neurons in the ARC) for each animal. All analyses were conducted on raw images without any post-acquisition modifications.

### Electrophysiological recordings in brain slices

*Calb1-Cre* mice injected with AAVs encoding Cre-dependent NpHR3.0-mCherry in the POA were anesthetized with isoflurane and transcardially perfused with ice-cold oxygenated (95% O_2_/5% CO_2_) high-sucrose solution (composition in mM: 2.5 KCl, 1.25 NaH_2_PO_4_, 2 Na_2_HPO_4_, 2 MgSO_4_, 213 sucrose, 26 NaHCO_3_). After brain dissection, coronal sections including the POA were cut at 250 μm using a vibratome (VT-1200S, Leica) in ice-cold oxygenated cutting solution. Brain slices were then incubated in artificial cerebrospinal fluid (ACSF; composition in mM: 126 NaCl, 2.5 KCl, 1.25 NaH_2_PO_4_, 1.25 Na_2_HPO_4_, 2 MgSO_4_, 10 glucose, 26 NaHCO_3_, 2 CaCl_2_) at 34 °C for at least 1 hour. Cells were identified under a fluorescent microscope and visualized by infrared differential interference contrast (BX51, Olympus). Whole-cell recordings were conducted using a MultiClamp700B amplifier and Digidata 1440A interface (Molecular Devices). Recording electrodes (3–5 MΩ, Borosilicate Glass, Sutter Instrument) were prepared with a micropipette puller (Sutter Instrument, model P97). The patch-clamp electrode was backfilled with an intracellular solution (composition in mM: 120 K-gluconate, 4 KCl, 10 HEPES, 10 sodium phosphocreatine, 4 Mg-ATP, and 0.3 Na_3_-GTP, pH 7.3, 265 mOsm). For optogenetic inhibition, stable action potentials were induced in NpHR-expressing neurons via current injection under current clamp, followed by repeated application of continuous yellow light onto the slices. Data were recorded using Clampe× 10.2 (Molecular Devices) and low-pass filtered at 10 kHz and sampled at 10 kHz under current clamp. All experiments were performed at 33 °C using a temperature controller (Warner, TC324B).

### Statistics

Statistical analyses were performed using Prism 10 (GraphPad Software). For bar graphs, values are presented as mean ± SEM. To compare the two groups, we first tested the data with the Lilliefors’ goodness of normality test; then variances were tested with F-test; if the data were fitted into normal distribution and equal variances, p values were calculated with paired or unpaired Student’s t-test. For unpaired data and not fitting a normal distribution, a non-parametric Mann–Whitney U-test was used. For data without equal variances, the paired or unpaired t test with Welch’s correction was used. For comparisons among data from paired three groups in Figure2h and Extended Figure5, repeated measures ANOVA followed by Šídák’s multiple comparisons test was used. For comparisons among data with the hypothesized value of the mean in Figure4, one sample test was used. Chi-square test with the p value adjusted for false discovery rate (FDR) of 1% using two-stage step-up method of Benjamini, Krieger and Yekutieli was used to analyze categorical data in Figure1e. FDR adjustment was also used in Extended Figure7 for correction of p-values after multiplied comparisons. For comparisons of survival distributions between two groups, Log-rank (Mantel-Cox) test was used. *p < 0.05, **p < 0.01, ***p < 0.001, ****p < 0.0001. For more detailed information, refer to Extended Data Table 1.

## Acknowledgement

We thank members of the Xu Lab for comments on the manuscript. This work was supported by the STI 2030-Major Projects (2021ZD0203203, 2021ZD02044400), the Strategic Priority Research Program of the Chinese Academy of Sciences (XDB32000000), the National Natural Science Foundation of China (Grant # 31871066, 31922028, 31900721, 32100817), and the Shanghai Municipal Science and Technology Major Project (2018 SHZDZX05). China Postdoctoral Science Foundation (Grant No. 2020TQ033 and 2020M681416).

## Author contributions

X.H.X. conceived the study; X.Z., Z.L.J. and S.S.L. performed the experiments, data analysis and figure plotting; X.Y.Liu, Y.Z.S and G.M.T performed stereotactic injections, CPP related behavioral test, and post-hoc histology; X.J.D performed RNAscope staining and related imaging; A.X.C., X.Y.Li and S.C.G., helped photometry recording and behavioral experiments; J.K.L helped in snRNAseq data analysis; X.Z.C. and W.Z. prepared samples for snRNASeq; Y.L.Z. helped maintain the colony; R.R.Y and X.C provided the rabies-SADδG -GFP viruses; X.Z. and X.H.X wrote the manuscript.

## Declaration of generative AI and AI-assisted technologies in the writing process

During the preparation of this work, Xiaohong Xu used ChatGTP/OpenAI in order to improve the readability of the text. After using this tool/service, the authors reviewed and edited the content as needed and take full responsibility for the content of the publication.

---

**Extended Data Figure 1.**
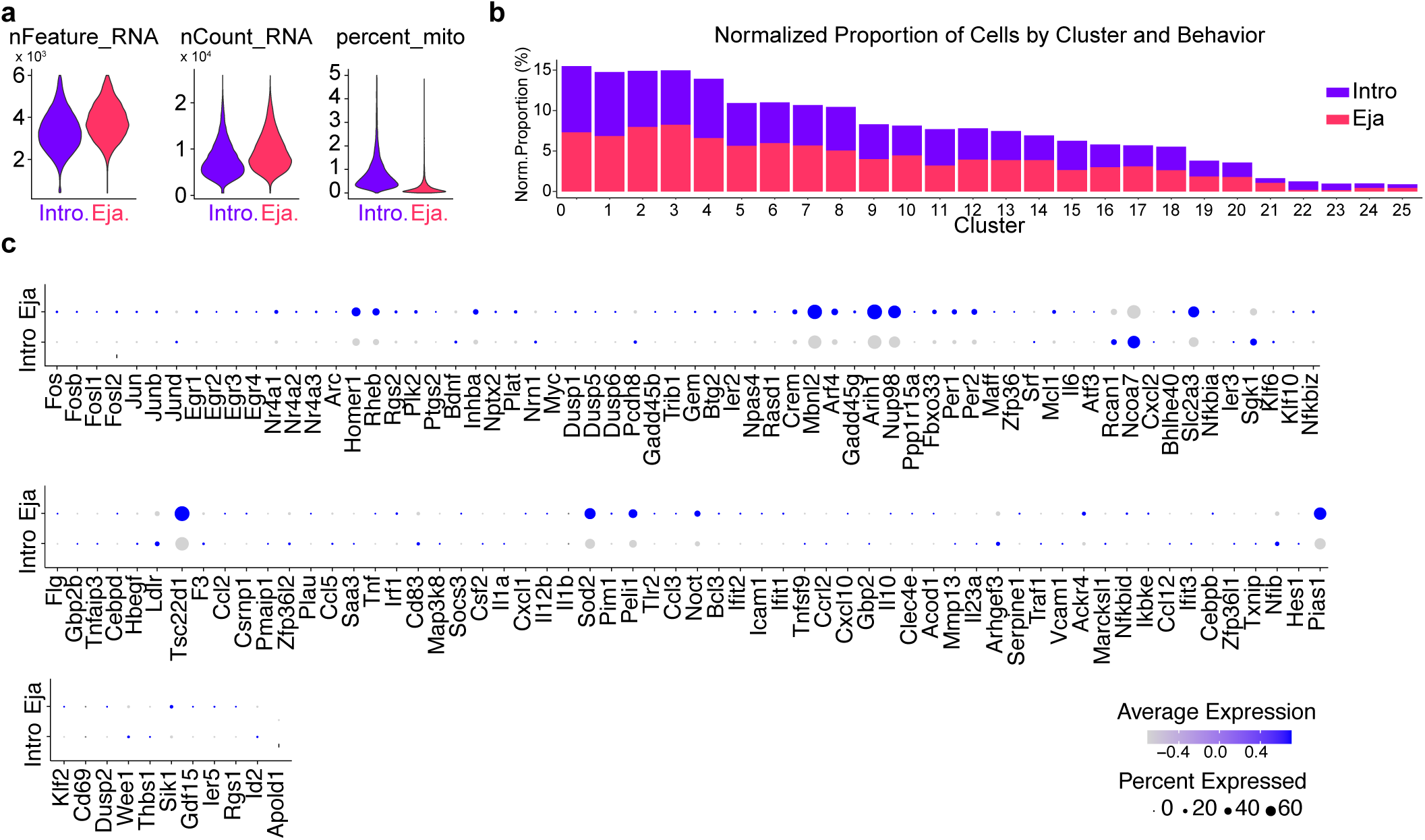
Increased IEG expression in POA neurons following ejaculation. **a.** Comparable quality of single-nucleus RNA-sequence data between cells from the intromission and ejaculation conditions, as demonstrated by similar number of detected genes (nFeature_RNA), total detected molecules (nCount_RNA), and similar percentage of mitochondria gene counts detected (percent_mito). **b.** Percentage of cells from the intromission (intro, blue) or the ejaculation (eja, red) conditions distributed across each POA neuron clusters. **c.** A heatmap illustrates the differential expression of IEGs (columns) between the ejaculation (top) and intromission (bottom) conditions, aggregating all POA neurons sequenced under each condition. The size of the circles represents the percentage of POA neurons expressing a given IEG, while the color intensity within the dots indicates the relative expression levels of the IEG.

**Extended Data Figure 2.**
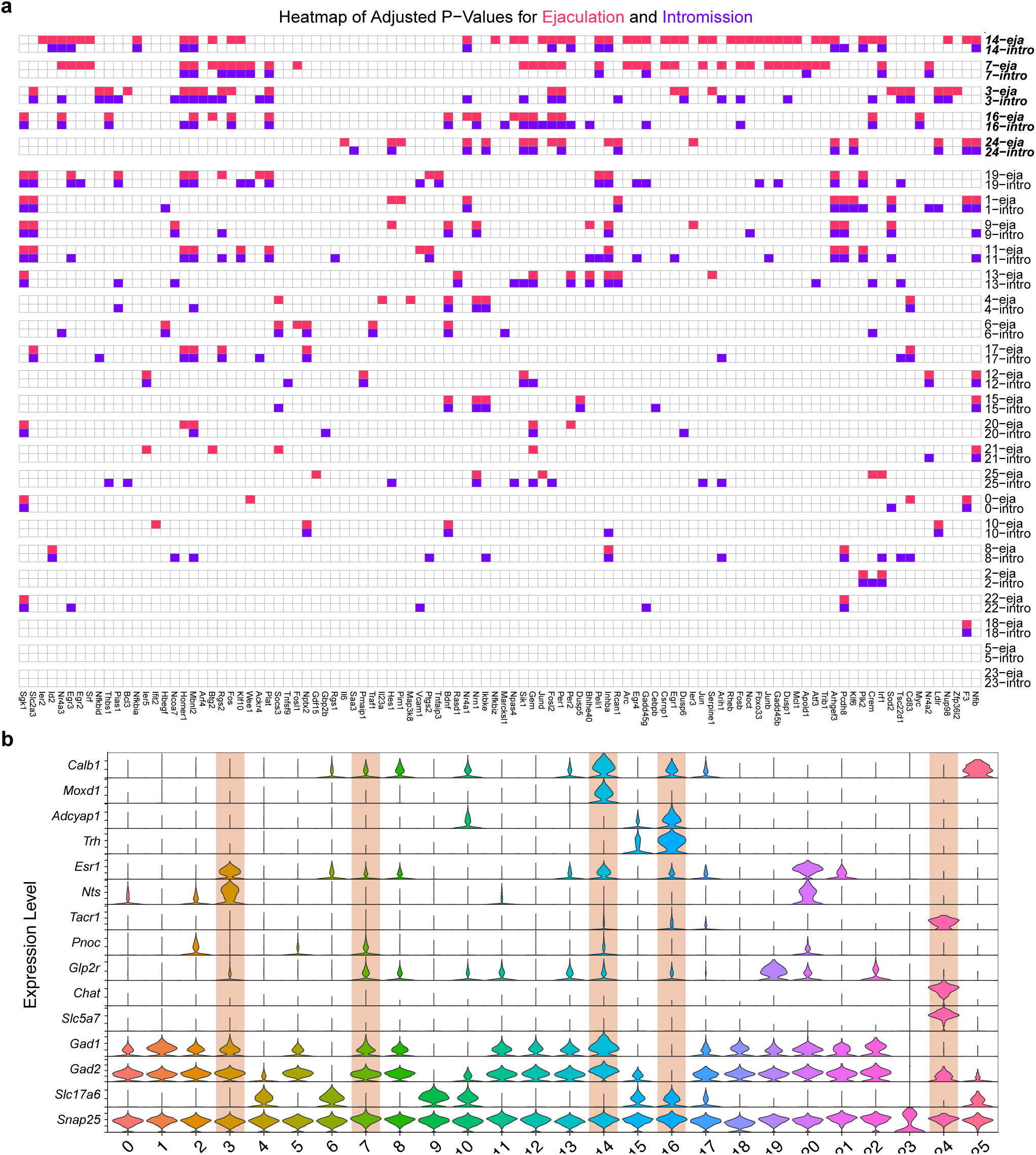
Activation of specific POA neuron clusters during ejaculation versus intromission. **a.** Filled blocks indicate the corresponding IEGs (columns) showed significant enrichment (adjusted p < 0.05) for a cluster (rows) under the ejaculation (red) or intromission (blue) condition. Clusters are ordered by the number of IEGs that showed enrichment under the ejaculation condition. Clusters #14, #7, #3, #16, #24 (in bold) show significant enrichment of more than 15 IEGs during ejaculation. **b.** Violin plots displaying the relative expression levels of marker genes (listed on the left) across the 26 neuron clusters, highlighting cluster #3 as *Esr1+/Nts+*, cluster #16 as *Calb1+/Adcyap1+*, and cluster #24 as *Tacr1+*.

**Extended Data Figure 3.**
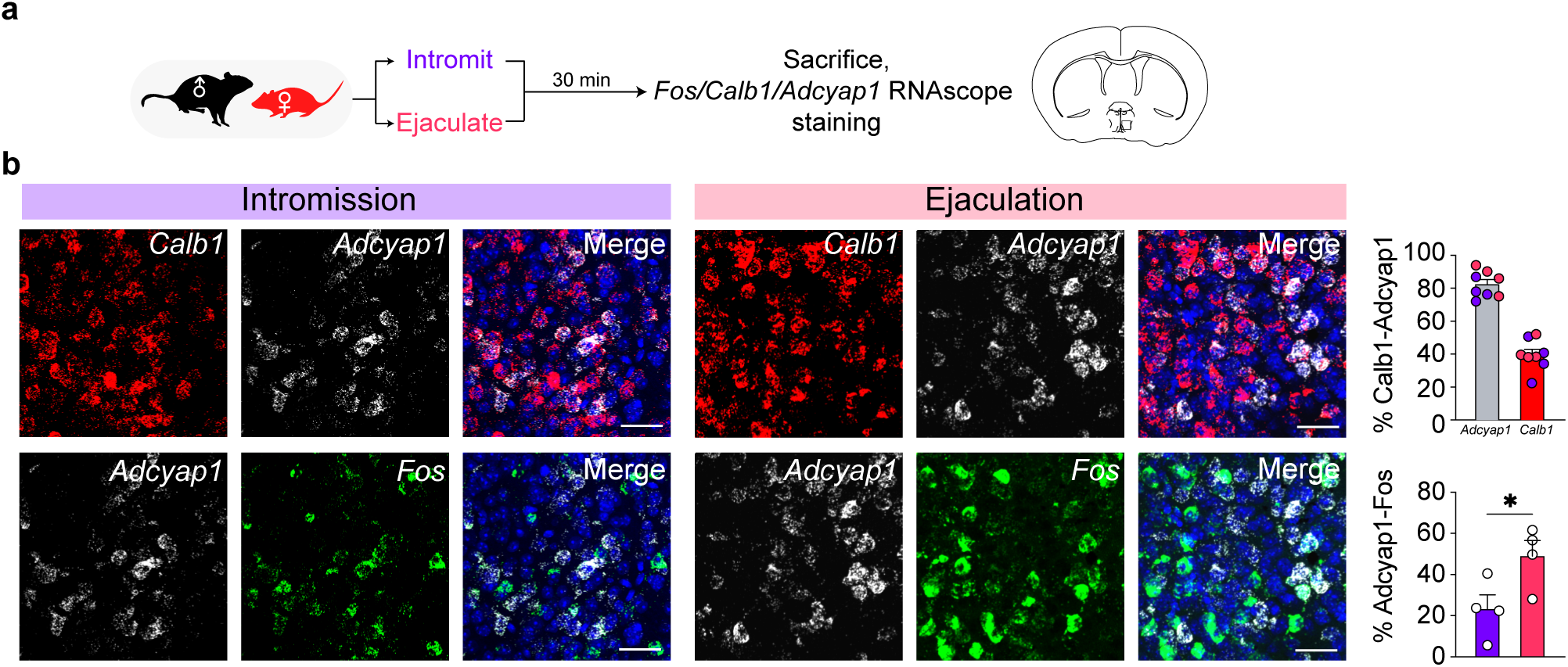
A subset of POA *Calb1+* neurons co-express Adcyap1 and are activated following ejaculation. **a.** Schematics of the experimental procedure. C57BL/6 male mice underwent a mating test, during which they either achieved ejaculation or intromission. Thirty minutes after the test, the mice were sacrificed, and their brains were processed for fluorescent *in situ* hybridization to detect *Calb1*, *Adcyap1* and *Fos* signals using RNAscope kits. **b.** Representative images (left) and quantification (right) are shown. The top row illustrates the overlap between *Calb1+* and *Adcyap1+* signals, with the color of the data points indicating the experimental group (blue for intromission and red for ejaculation). The bottom row displays the overlap between *Adcyap1+* and *Fos+* signals under intromission (left, blue) and ejaculation (right, red) conditions. Scale bar, 50 μm. n = 4 males/group. Values are presented as mean ± SEM. *p < 0.05.

**Extended Data Figure 4.**
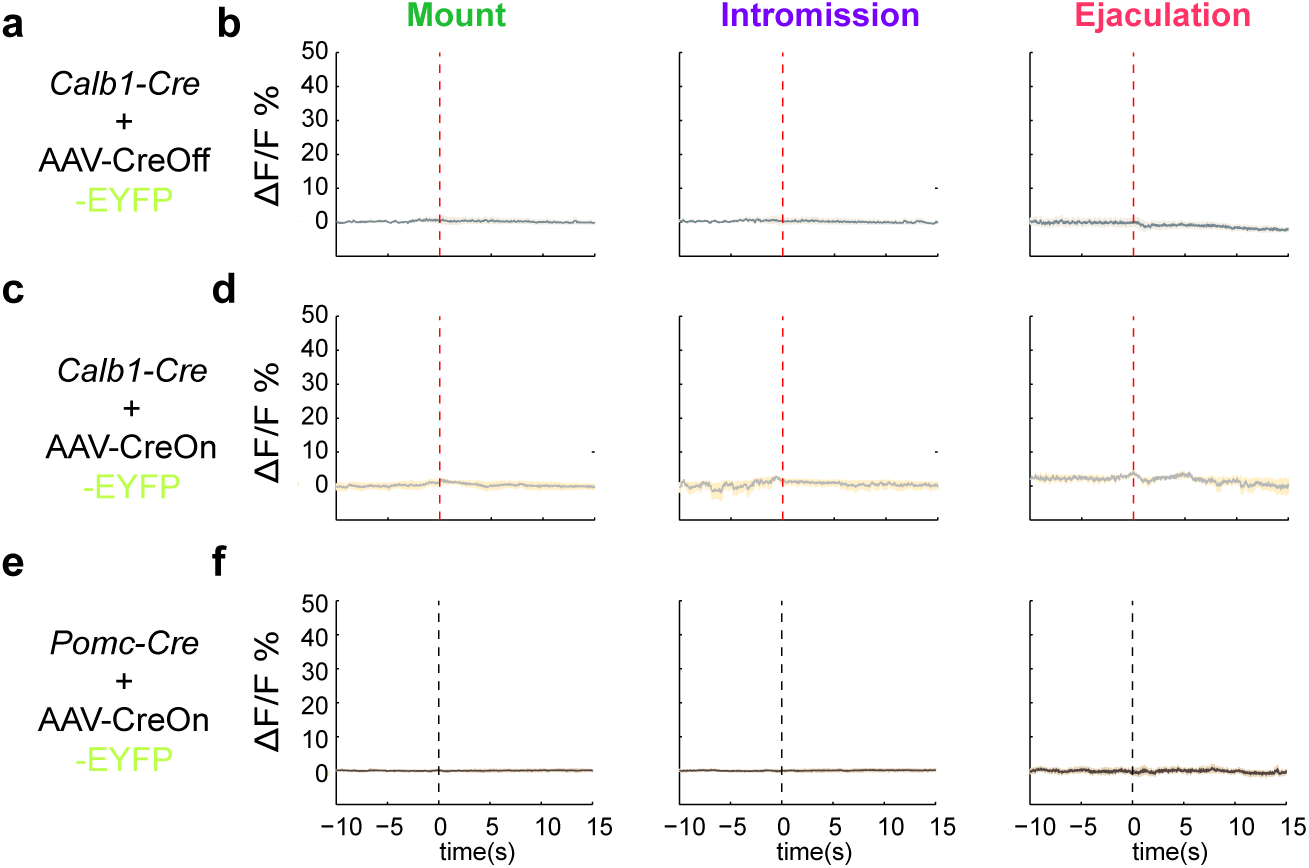
No changes in fluorescent signals were detected in EYFP-expressing neurons during male mating. Fluorescent signals were recorded via implanted optic fibers during male mating in *Calb1-Cre* male mice injected with AAVs encoding Cre-Off or Cre-On EYFP in the POA to label *Calb1+* (**a,** top row) and *non-Calb1* neurons (**c**, middle row), or in *Pomc-Cre* male mice (C57BL/6 background) injected with AAVs encoding Cre-On EYFP in the ARC to label *Pomc+* neurons (**e,** bottom row). ΔF/F values corresponding to the indicated behaviors (color coded, labelled on the top) were plotted accordingly (**b, d, f**). n = 5 males for *Calb1+* group, 6 males for *non-Calb1* group, and 3 males for *Pomc+* group.

**Extended Data Figure 5.**
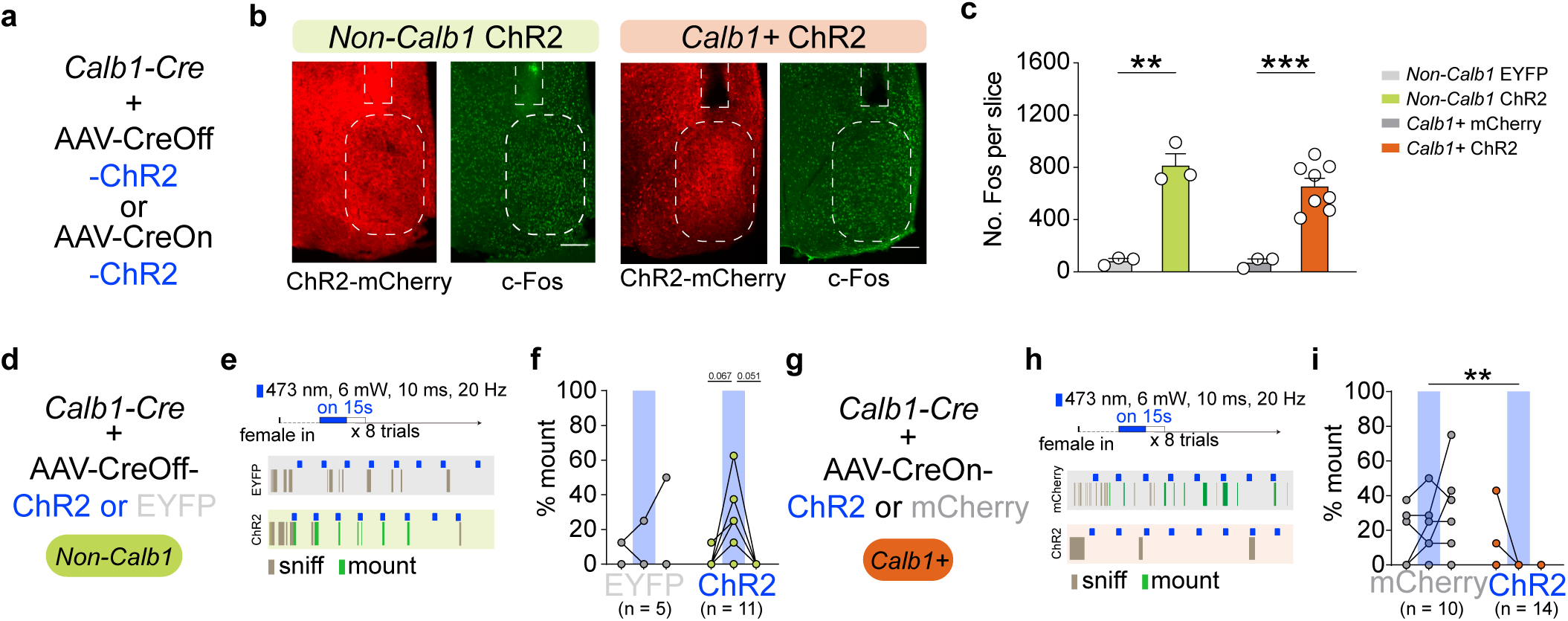
Optogenetic activation of POA *Calb1+* and *non-Calb1* neurons. **a.** Schematic representation of the experimental design: AAVs encoding Cre-Off or Cre-On ChR2-mCherry were injected unilaterally into the POA of *Calb1-Cre* male mice to label *non-Calb1* and *Calb1+* neurons, respectively. **b.** Representative images showing the POA labeling with implanted optic fiber, and light-induced c-Fos expression in *non-Calb1* or *Calb1+* neuron-targeted group. Scale bar, 200 μm. **c.** Quantification of POA c-Fos signals following light stimulation in the respective groups. n = 3 males for *non-Calb1* EYFP group, 3 males for *non-Calb1* ChR2 group, 3 males for *Calb1*+ mCherry group, and 8 males for *Calb1*+ ChR2 group. **d.-i.** Optogenetic activation of POA *non-Calb1* and *Calb1*+ neurons differentially affected mount. **d, g**: Diagrams of the viral strategy; **e, h**: Light delivery protocol and raster plots of behavior for representative trials in both experimental and control groups. **f, i**: Quantification of the percentage of light delivery period (denoted by blue bars) where light-activated mounting behavior occurred. Activation of *non-Calb1* neurons promotes mounting behavior (**d-f**), while activation of *Calb1*+ neurons inhibits mounting behavior (**g-i**). n = 11 males for the *non-Calb1* ChR2 group, 5 males for the *non-Calb1* EYFP group, 14 males for the *Calb1+* ChR2 group, 10 males for the *Calb1+* mCherry group. Values are presented as mean ± SEM. **p < 0.01, ***p < 0.001.

**Extended Data Figure 6.**
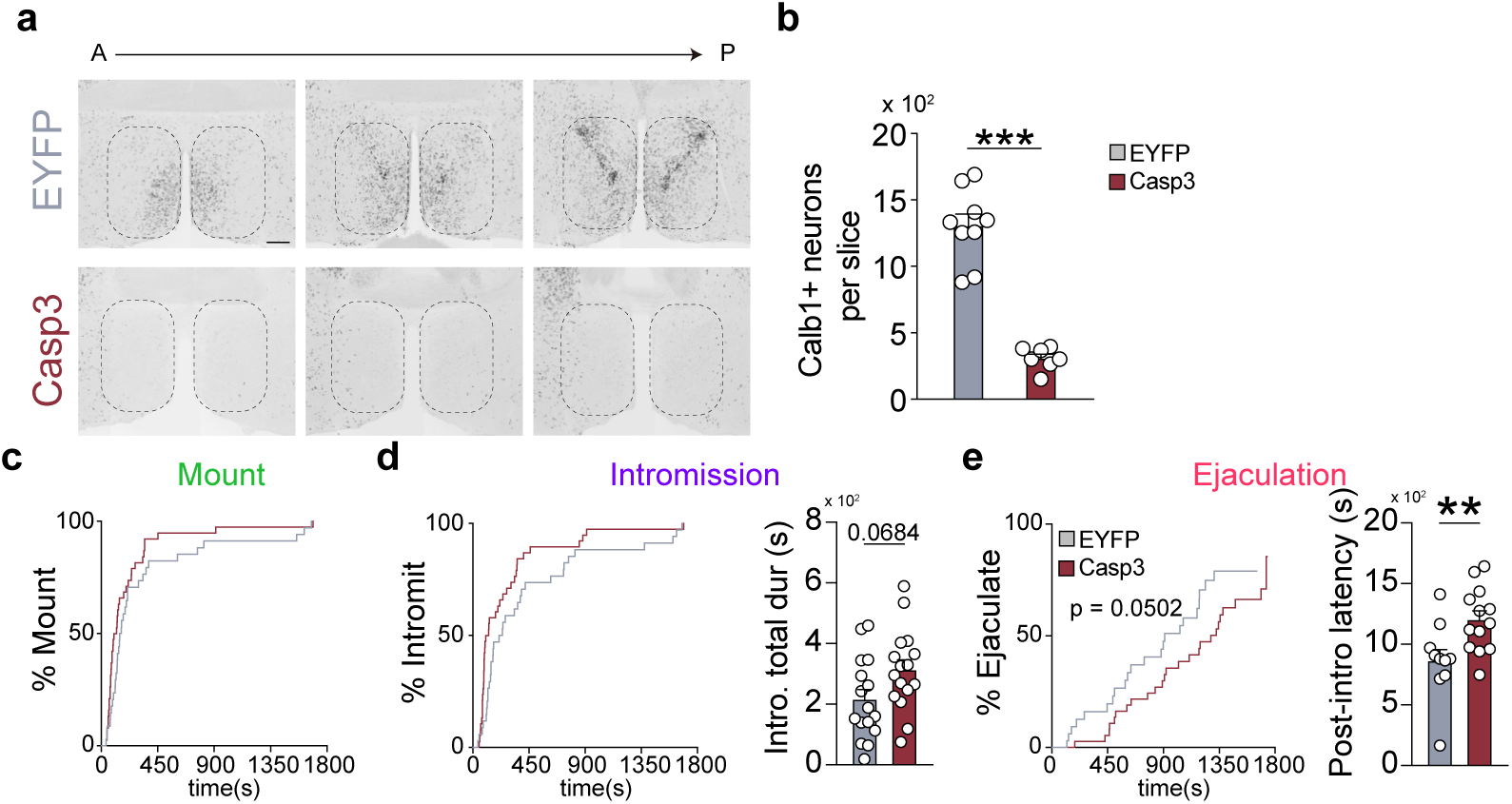
Ablation of POA *Calb1+* neurons delays ejaculation but does not affect the initiation of mounting or intromission during male mating. **a.** Serial montage of representative images from anterior (A) to posterior (P) showing the elimination of *Calb1+ in situ* signals in the POA (outlined) after AAVs encoding Cre-On Caspase3 (Casp3) were injected into the POA of *Calb1-Cre* male mice to ablate *Calb1+* neurons. Control males were injected with AAVs encoding Cre-On EYFP. Scale bar, 200 μm. **b.** Quantification of the number of POA *Calb1+* neurons in control (EYFP-injected) and Casp3-injected males. n = 7 Casp3 males and 9 EYFP males. **c.-e.** Casp3-injected males were comparable to control males in the initiation of mounting (**c**) and intromission (**d**) during male mating tests but the total duration of intromission was extended in the Casp3 group. Moreover, the percentage of males that ejaculated at a given time after introduction of a female (time ‘0’) was reduced in Casp3-injected males and the latency to ejaculation following the first intromission (Post-intro latency) was significantly increased. n = 17 Casp3 and 16 EYFP males. Values are presented as mean ± SEM. **p < 0.01, ***p < 0.001.

**Extended Data Figure 7.**
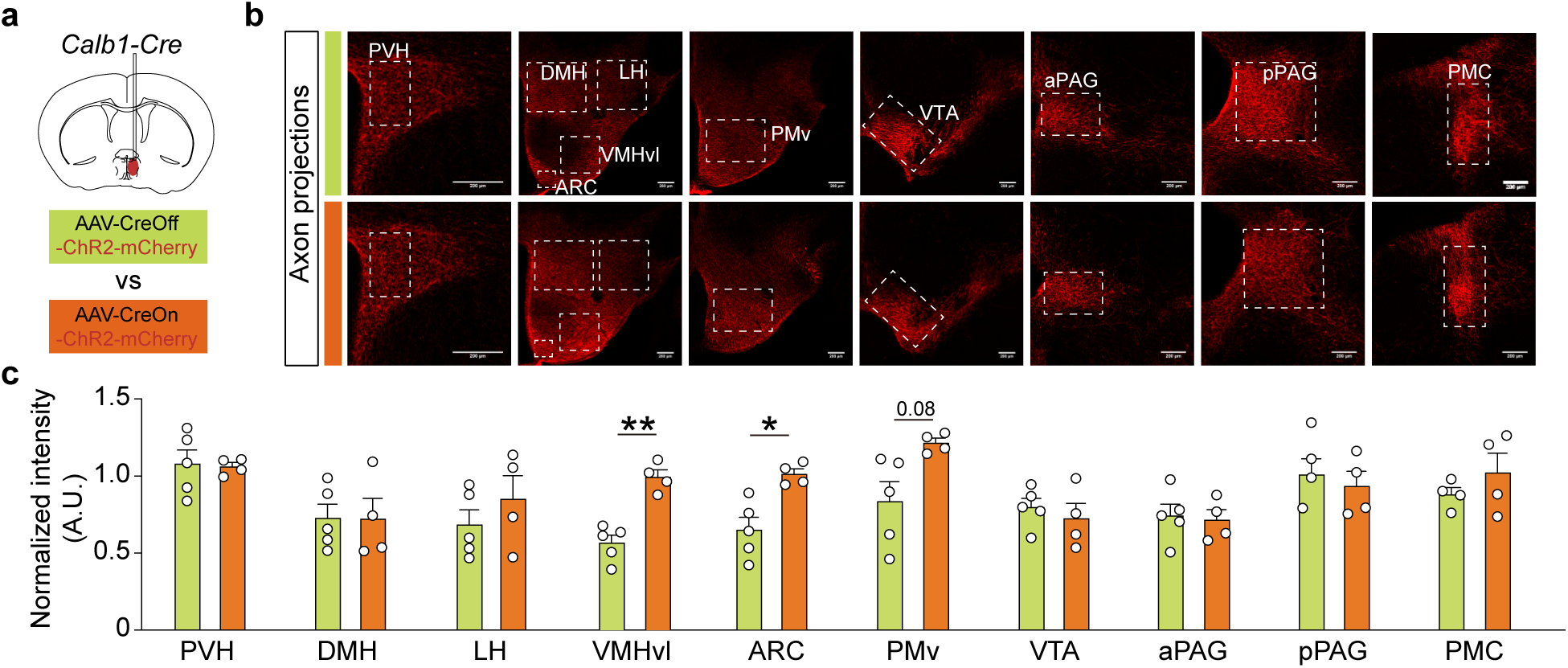
Denser innervation of ARC and the ventrolateral division of VMH (VMHvl) by POA *Calb1+* neurons. **a.** Schematic representation of the experimental procedure: AAVs encoding Cre-Off or Cre-On ChR2-mCherry were injected into the POA of *Calb1-Cre* male mice to label the axon terminals of *non-Calb1* and *Calb1+* neurons respectively. Scale bar, 200 μm. **b.-c.** Representative images (**b**) and quantification and comparison of axon projection strength of POA *non-Calb1* (top row, green) and *Calb1+* (bottom row, red) neurons in indicated brain regions. n = 4 *Calb1+* males and 5 *non-Calb1* males. Values are presented as mean ± SEM. * adjusted p < 0.05, ** adjusted p < 0.01.

**Extended Data Figure 8.**
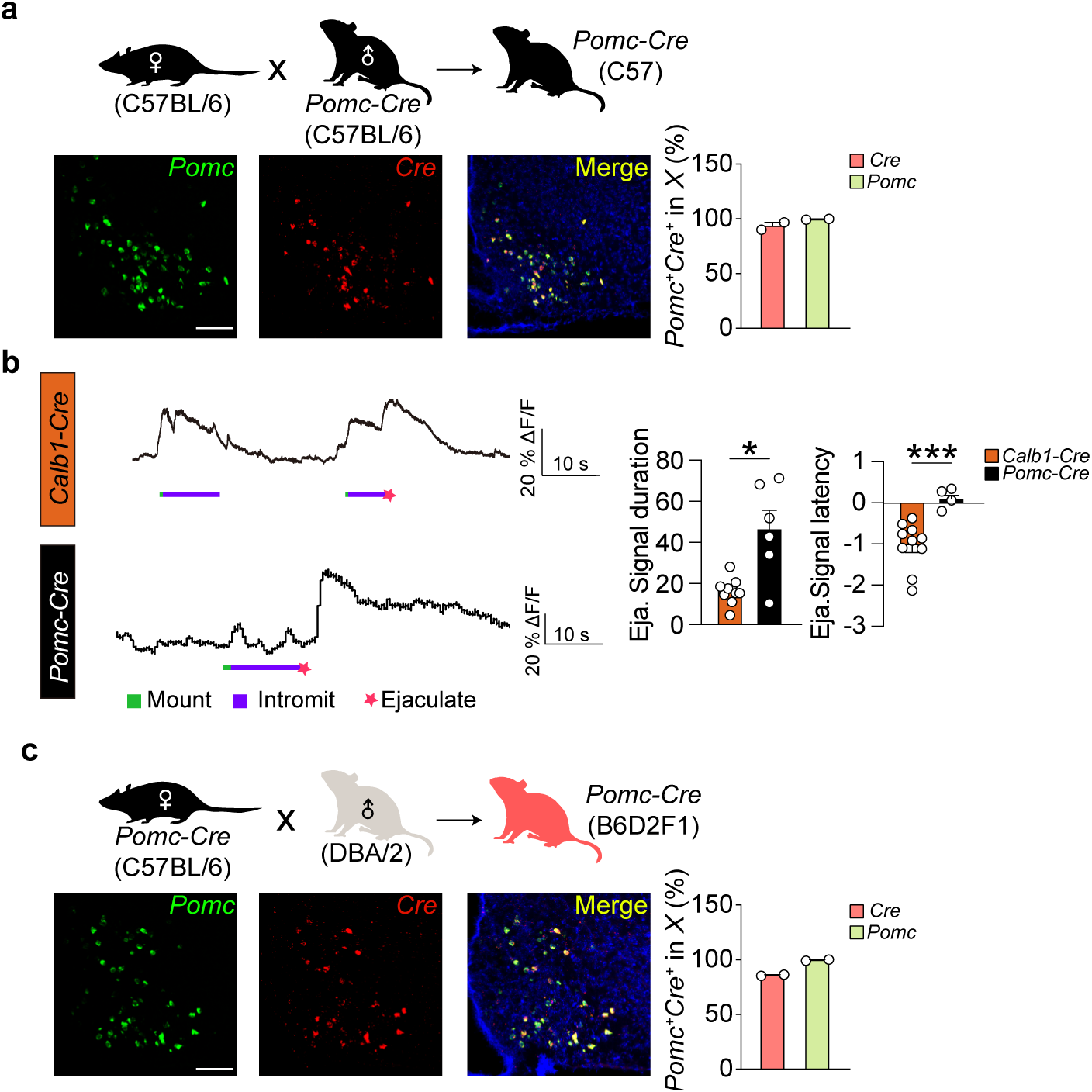
Validation of the *Pomc-Cre* line and comparison of ejaculation-associated activity in *Pomc+* and *Calb1+* neurons. **a, c.** Illustration of the *Pomc-Cre* line maintained on the C57BL/6 background (**a**) and the B6D2F1 hybrid background (**c**). Representative RNAscope fluorescent *in situ* hybridization images with the quantification show overlap between *Cre* expression and *Pomc* signals. Scale bar, 50 μm. n = 2/group. **b.** Representative traces of GCamp6s signals (ΔF/F) recorded in POA *Calb1+* neurons (top) and ARC *Pomc+* neurons (bottom). GCamp6s signal duration (left) during ejaculation (Eja.) and latency relative to ejaculation as time ‘0’ (right) were plotted. The quantification showing delayed but more sustained activation of *Pomc+* neurons during ejaculation compared to POA *Calb1+* neurons. n = 10 *Calb1+* males and 6 *Pomc+* males. Values are presented as mean ± SEM. *p < 0.05 ***p < 0.001.

**Extended Data Figure 9.**
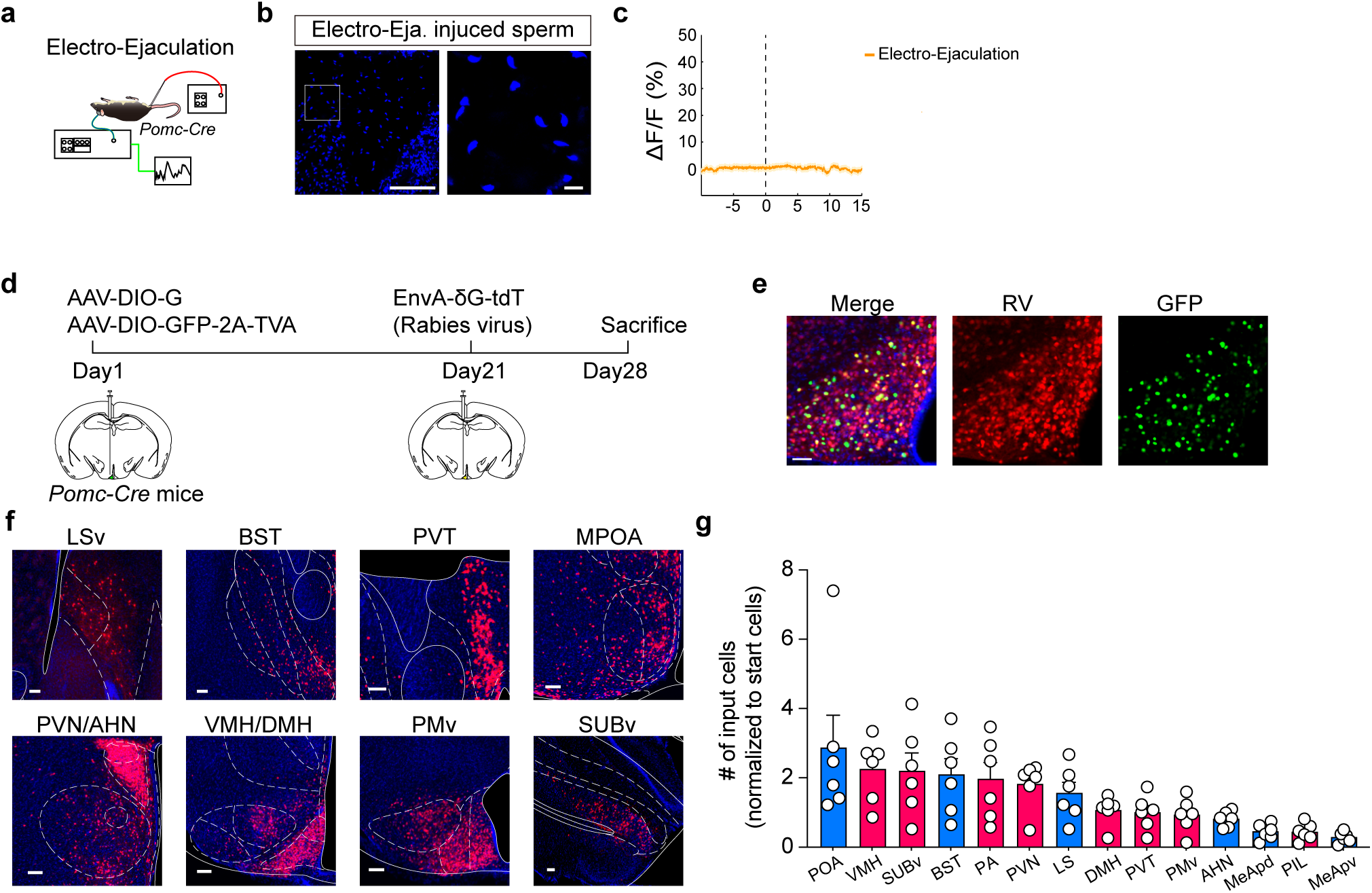
Ejaculation induced by electrical currents through a rectum probe did not increase activity in *Pomc+* neurons, and brain regions that provide inputs to ARC *Pomc+* neurons. **a.** Schematics of the experimental setup. **b.** Histological examination of sperm in the ejaculates triggered with electrical currents. The right image zooms in on the white box from the left image. Scale bar, 100 μm for the left image and 10 μm for the right image. **c.** Fiber photometry recordings of *Pomc* GCaMP6s signals show no changes in fluorescent activity during electrically induced ejaculation (time “0”). n = 2. **d.** Schematics of the viral strategy using pseudo-rabies virus-mediated retrograde tracing from ARC *Pomc+* neurons. **e.-f.** Representative images showing starter cells in ARC (**e**, scale bar, 50 μm) and labeled upstream neurons in the indicated brain regions (**f**, scale bar, 100 μm). **g.** Quantification of inputs from various inhibitory (blue) or excitatory (red) brain regions. n = 6.

**Extended Data Video 1-4.** Video clips played at 2x speed showing male mating behaviors at the top, with mount (green bar), intromission (red bar), and ejaculation (blue bar) annotated on the video, alongside recordings of GCaMP6s signals from defined neuron populations: 1. POA *Calb1+* neurons; 2. POA *non-Calb1* neurons; 3. ARC *Pomc+* neurons on the C57BL/6 background; 4. ARC *Pomc+* neurons on the B6D2F1 hybrid background.

**Extended Data Table 1.**
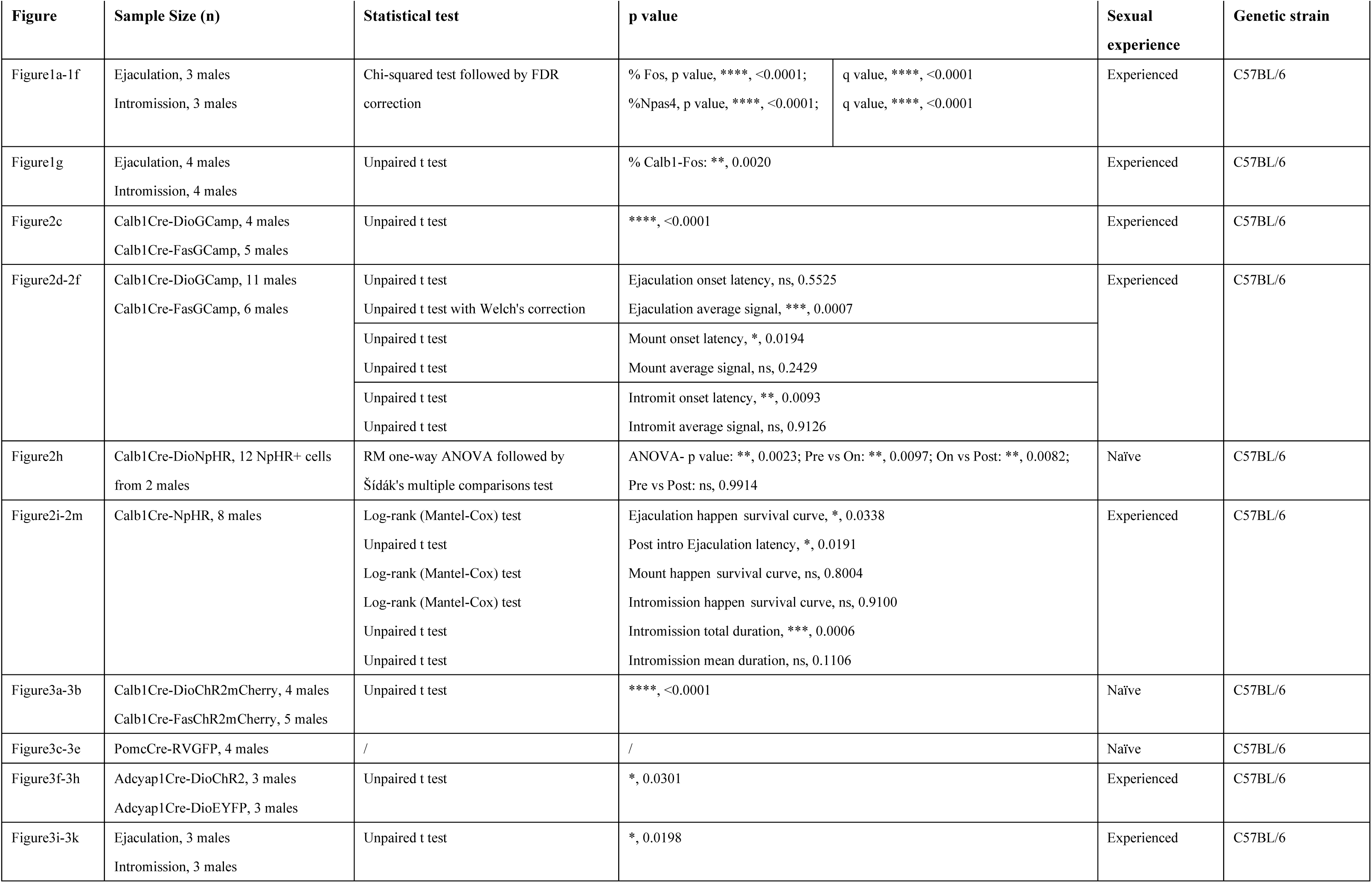
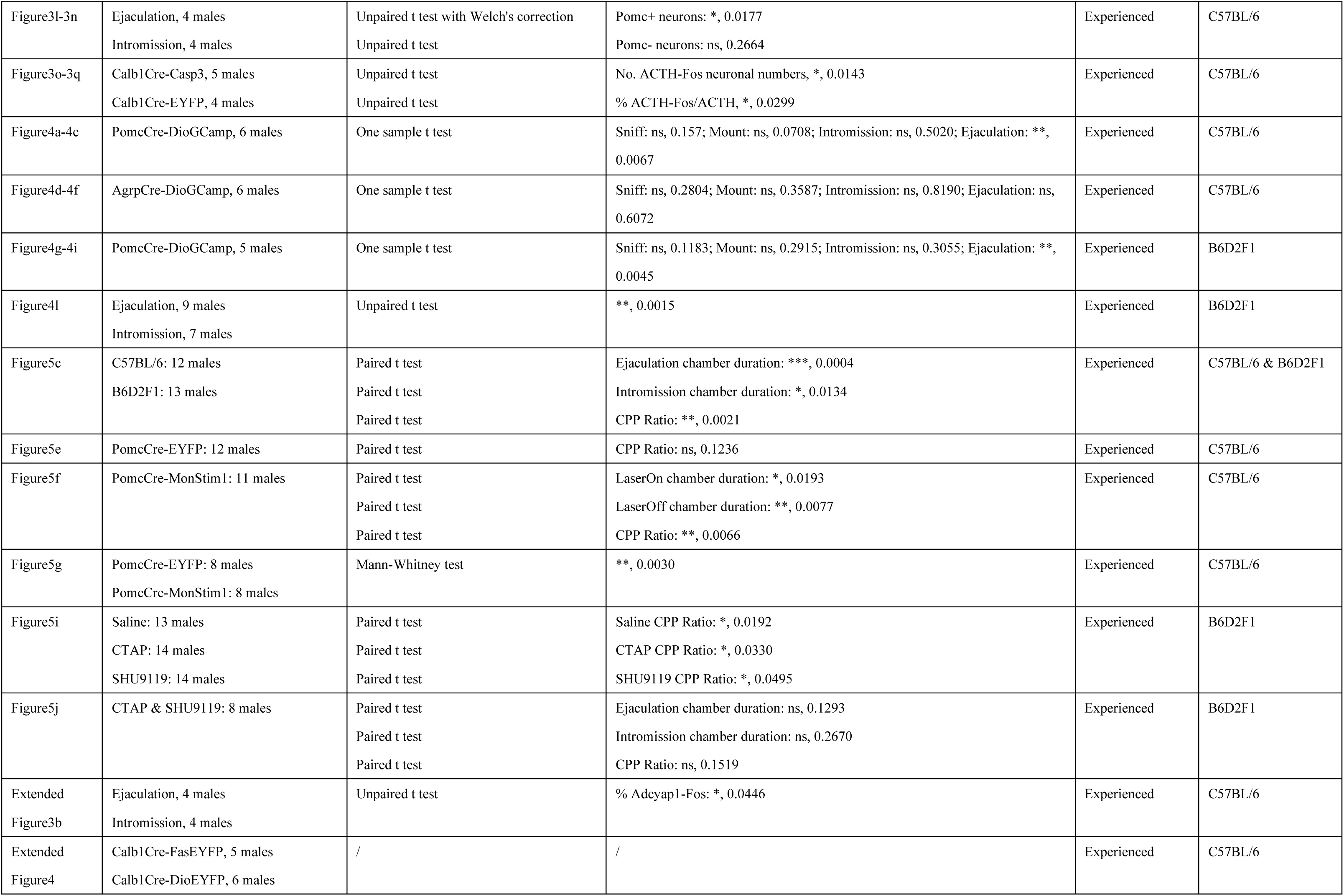
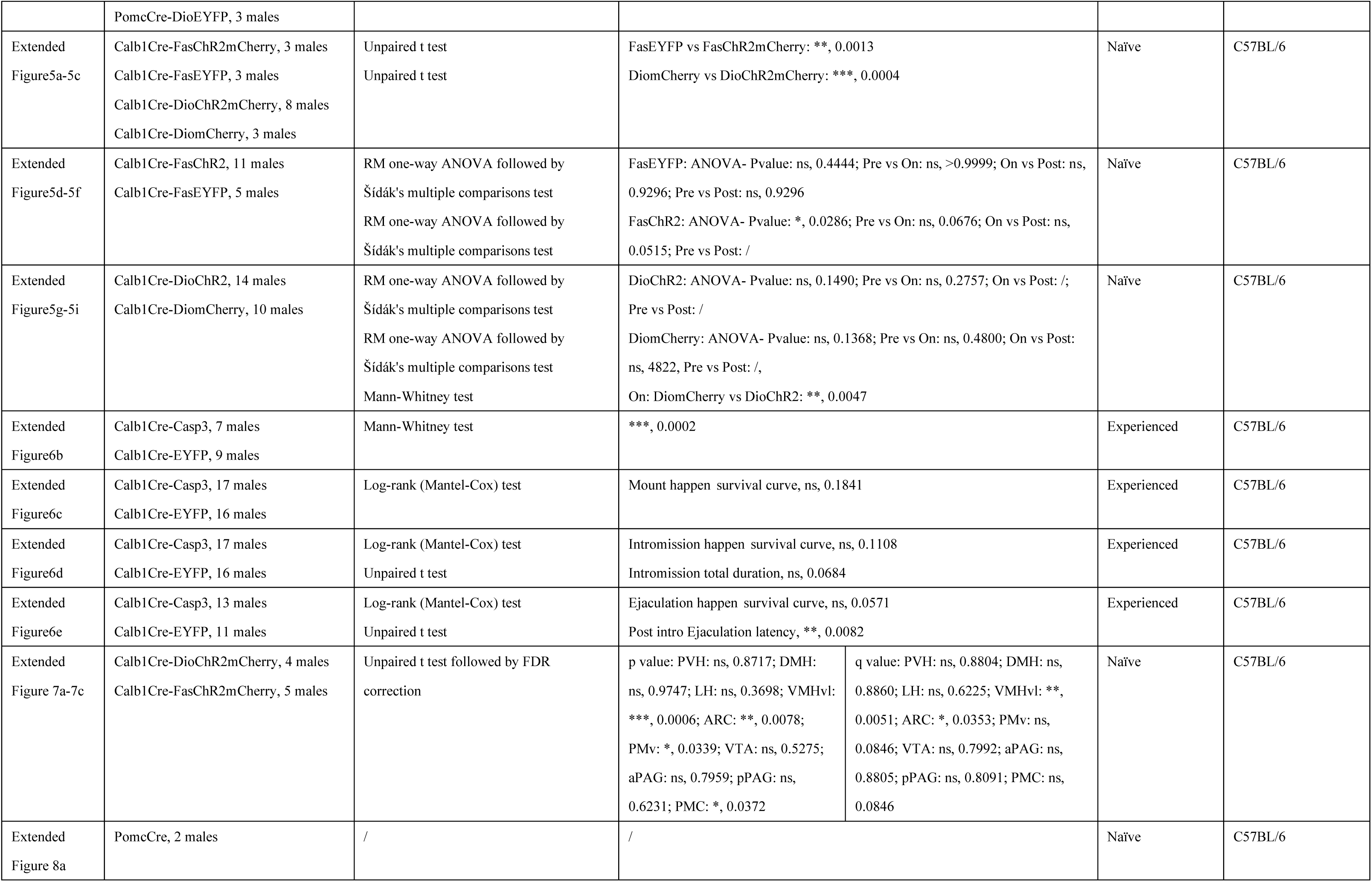
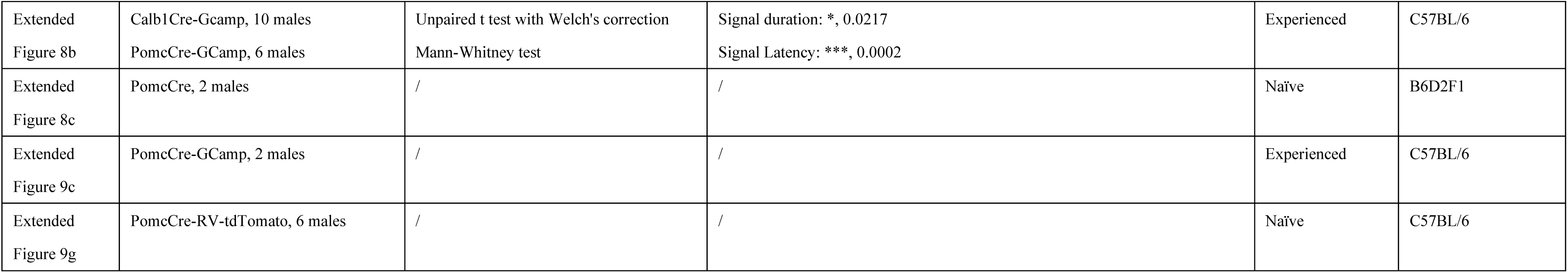
This table provided detailed statistical information, along with the sexual experience and genetic background of the males, corresponding to each figure panel.

## Data Availability

All unique/stable reagents generated in this study are available from the Lead Contact without restriction.

## Reference

1 Perez, M., Hoppe, J., Landau, A. M. & Winterdahl, M. Development and validation of the Male Post-coital Affect Scale for heterosexual men. Acta Neuropsychiatr 35, 241–246 (2023). 10.1017/neu.2023.12

2 Mah, K. & Binik, Y. M. Do all orgasms feel alike? Evaluating a two-dimensional model of the orgasm experience across gender and sexual context. J Sex Res 39, 104–113 (2002). 10.1080/00224490209552129

3 Pfaus, J. G. et al. Do rats have orgasms? Socioaffect Neurosci Psychol 6, 31883 (2016). 10.3402/snp.v6.31883

4 Yu, Z. X., Zha, X. & Xu, X. H. Estrogen-responsive neural circuits governing male and female mating behavior in mice. Curr Opin Neurobiol 81, 102749 (2023). 10.1016/j.conb.2023.102749

5 Harno, E., Gali Ramamoorthy, T., Coll, A. P. & White, A. POMC: The Physiological Power of Hormone Processing. Physiol Rev 98, 2381–2430 (2018). 10.1152/physrev.00024.2017

6 Xiao, W. et al. Neural circuit control of innate behaviors. Sci China Life Sci 65, 466–499 (2022). 10.1007/s11427-021-2043-2

7 Hashikawa, K., Hashikawa, Y., Falkner, A. & Lin, D. The neural circuits of mating and fighting in male mice. Curr Opin Neurobiol 38, 27–37 (2016). 10.1016/j.conb.2016.01.006

8 Wei, Y. C. et al. Medial preoptic area in mice is capable of mediating sexually dimorphic behaviors regardless of gender. Nat Commun 9, 279 (2018). 10.1038/s41467-017-02648-0

9 Bayless, D. W. et al. A neural circuit for male sexual behavior and reward. Cell 186, 3862–3881 e3828 (2023). 10.1016/j.cell.2023.07.021

10 Jiao, Z. L. et al. Acute Recruitment of VTA Dopamine Neurons by mPOA Esr1+ Neurons to Facilitate Consummatory Male Mating Actions. Neurosci Bull (2024). 10.1007/s12264-024-01288-x

11 Dai, B. et al. Responses and functions of dopamine in nucleus accumbens core during social behaviors. Cell Rep 40, 111246 (2022). 10.1016/j.celrep.2022.111246

12 Qian, T. et al. A genetically encoded sensor measures temporal oxytocin release from different neuronal compartments. Nat Biotechnol 41, 944–957 (2023). 10.1038/s41587-022-01561-2

13 Alwaal, A., Breyer, B. N. & Lue, T. F. Normal male sexual function: emphasis on orgasm and ejaculation. Fertil Steril 104, 1051–1060 (2015). 10.1016/j.fertnstert.2015.08.033

14 Clement, P. & Giuliano, F. Physiology and Pharmacology of Ejaculation. Basic Clin Pharmacol Toxicol 119 **Suppl 3**, 18–25 (2016). 10.1111/bcpt.12546

15 Corty, E. W. Perceived ejaculatory latency and pleasure in different outlets. J Sex Med 5, 2694–2702 (2008). 10.1111/j.1743-6109.2008.00939.x

16 Truitt, W. A. & Coolen, L. M. Identification of a potential ejaculation generator in the spinal cord. Science 297, 1566–1569 (2002). 10.1126/science.1073885

17 Coolen, L. M., Allard, J., Truitt, W. A. & McKenna, K. E. Central regulation of ejaculation. Physiol Behav 83, 203–215 (2004). 10.1016/j.physbeh.2004.08.023

18 Veening, J. G. & Coolen, L. M. Neural mechanisms of sexual behavior in the male rat: emphasis on ejaculation-related circuits. Pharmacol Biochem Behav 121, 170–183 (2014). 10.1016/j.pbb.2013.12.017

19 Seecof, R. & Tennant, F. S., Jr. Subjective perceptions to the intravenous “rush” of heroin and cocaine in opioid addicts. Am J Drug Alcohol Abuse 12, 79–87 (1986). 10.3109/00952998609083744

20 Zhou, X. et al. Hyperexcited limbic neurons represent sexual satiety and reduce mating motivation. Science 379, 820–825 (2023). 10.1126/science.abl4038

21 Zhang, S. X. et al. Hypothalamic dopamine neurons motivate mating through persistent cAMP signalling. Nature 597, 245–249 (2021). 10.1038/s41586-021-03845-0

22 Baum, M. J. & Everitt, B. J. Increased expression of c-fos in the medial preoptic area after mating in male rats: role of afferent inputs from the medial amygdala and midbrain central tegmental field. Neuroscience 50, 627–646 (1992). 10.1016/0306-4522(92)90452-8

23 Kollack-Walker, S. & Newman, S. W. Mating-induced expression of c-fos in the male Syrian hamster brain: role of experience, pheromones, and ejaculations. J Neurobiol 32, 481–501 (1997). 10.1002/(sici)1097-4695(199705)32:5<481::aid-neu4>3.0.co;2-1

24 Coolen, L. M., Peters, H. J. & Veening, J. G. Fos immunoreactivity in the rat brain following consummatory elements of sexual behavior: a sex comparison. Brain Res 738, 67–82 (1996). 10.1016/0006-8993(96)00763-9

25 Wu, Y. E., Pan, L., Zuo, Y., Li, X. & Hong, W. Detecting Activated Cell Populations Using Single-Cell RNA-Seq. Neuron 96, 313–329 e316 (2017). 10.1016/j.neuron.2017.09.026

26 Osterhout, J. A. et al. A preoptic neuronal population controls fever and appetite during sickness. Nature 606, 937–944 (2022). 10.1038/s41586-022-04793-z

27 Tsuneoka, Y. et al. Moxd1 Is a Marker for Sexual Dimorphism in the Medial Preoptic Area, Bed Nucleus of the Stria Terminalis and Medial Amygdala. Front Neuroanat 11, 26 (2017). 10.3389/fnana.2017.00026

28 Tsukahara, S. & Morishita, M. Sexually Dimorphic Formation of the Preoptic Area and the Bed Nucleus of the Stria Terminalis by Neuroestrogens. Front Neurosci 14, 797 (2020). 10.3389/fnins.2020.00797

29 Bloch, G. J. & Gorski, R. A. Cytoarchitectonic analysis of the SDN-POA of the intact and gonadectomized rat. J Comp Neurol 275, 604–612 (1988). 10.1002/cne.902750408

30 Arendash, G. W. & Gorski, R. A. Effects of discrete lesions of the sexually dimorphic nucleus of the preoptic area or other medial preoptic regions on the sexual behavior of male rats. Brain Res Bull 10, 147–154 (1983). 10.1016/0361-9230(83)90086-2

31 Gorski, R. A., Harlan, R. E., Jacobson, C. D., Shryne, J. E. & Southam, A. M. Evidence for the existence of a sexually dimorphic nucleus in the preoptic area of the rat. J Comp Neurol 193, 529–539 (1980). 10.1002/cne.901930214

32 Qi, L. et al. Krause corpuscles are genital vibrotactile sensors for sexual behaviours. Nature 630, 926–934 (2024). 10.1038/s41586-024-07528-4

33 Millington, G. W. The role of proopiomelanocortin (POMC) neurones in feeding behaviour. Nutr Metab (Lond) 4, 18 (2007). 10.1186/1743-7075-4-18

34 McGill, T. E. & Tucker, G. R. Genotype and Sex Drive in Intact and in Castrated Male Mice. Science 145, 514–515 (1964). 10.1126/science.145.3631.514

35 Roth-Deri, I., Green-Sadan, T. & Yadid, G. Beta-endorphin and drug-induced reward and reinforcement. Prog Neurobiol 86, 1–21 (2008). 10.1016/j.pneurobio.2008.06.003

36 Paredes, R. G. Evaluating the neurobiology of sexual reward. ILAR J 50, 15–27 (2009). 10.1093/ilar.50.1.15

37 Tenk, C. M., Wilson, H., Zhang, Q., Pitchers, K. K. & Coolen, L. M. Sexual reward in male rats: effects of sexual experience on conditioned place preferences associated with ejaculation and intromissions. Horm Behav 55, 93–97 (2009). 10.1016/j.yhbeh.2008.08.012

38 Kim, S. et al. Non-invasive optical control of endogenous Ca(2+) channels in awake mice. Nat Commun 11, 210 (2020). 10.1038/s41467-019-14005-4

39 Klawonn, A. M. et al. Motivational valence is determined by striatal melanocortin 4 receptors. J Clin Invest 128, 3160–3170 (2018). 10.1172/JCI97854

40 Haynes, W. G., Morgan, D. A., Djalali, A., Sivitz, W. I. & Mark, A. L. Interactions between the melanocortin system and leptin in control of sympathetic nerve traffic. Hypertension 33, 542–547 (1999). 10.1161/01.hyp.33.1.542

41 Wang, X. et al. A non-peptide substance P antagonist (CP-96,345) inhibits morphine-induced NF-kappa B promoter activation in human NT2-N neurons. J Neurosci Res 75, 544–553 (2004). 10.1002/jnr.10873

42 Coolen, L. M., Fitzgerald, M. E., Yu, L. & Lehman, M. N. Activation of mu opioid receptors in the medial preoptic area following copulation in male rats. Neuroscience 124, 11–21 (2004). 10.1016/j.neuroscience.2003.10.045

43 Pfaus, J. G. & Gorzalka, B. B. Opioids and sexual behavior. Neurosci Biobehav Rev 11, 1–34 (1987). 10.1016/s0149-7634(87)80002-7

44 Chessick, R. D. The “pharmacogenic orgasm” in the drug addict. Arch Gen Psychiatry 3, 545–556 (1960). 10.1001/archpsyc.1960.01710050095010

45 Toda, C., Santoro, A., Kim, J. D. & Diano, S. POMC Neurons: From Birth to Death. Annu Rev Physiol 79, 209–236 (2017). 10.1146/annurev-physiol-022516-034110

46 Kennedy, A. et al. Stimulus-specific hypothalamic encoding of a persistent defensive state. Nature 586, 730–734 (2020). 10.1038/s41586-020-2728-4

47 Cheung, K. Y. M., Nair, A., Li, L. Y., Shapiro, M. G. & Anderson, D. J. Population coding of predator imminence in the hypothalamus. bioRxiv (2024). 10.1101/2024.08.12.607651

48 Mercer, A. J., Hentges, S. T., Meshul, C. K. & Low, M. J. Unraveling the central proopiomelanocortin neural circuits. Front Neurosci 7, 19 (2013). 10.3389/fnins.2013.00019

49 Jiao, Z. et al. Projectome-defined subtypes and modular intra-hypothalamic subnetworks of peptidergic neurons. bioRxiv, 2023.2005.2025.542241 (2023). 10.1101/2023.05.25.542241

50 Mountoufaris, G. et al. A line attractor encoding a persistent internal state requires neuropeptide signaling. Cell (2024). 10.1016/j.cell.2024.08.015

51 Nair, A. et al. An approximate line attractor in the hypothalamus encodes an aggressive state. Cell 186, 178–193 e115 (2023). 10.1016/j.cell.2022.11.027

52 Vinograd, A., Nair, A., Kim, J., Linderman, S. W. & Anderson, D. J. Causal evidence of a line attractor encoding an affective state. Nature (2024). 10.1038/s41586-024-07915-x

53 Liu, M., Nair, A., Coria, N., Linderman, S. W. & Anderson, D. J. Encoding of female mating dynamics by a hypothalamic line attractor. Nature (2024). 10.1038/s41586-024-07916-w

54 Yang, T. et al. Hypothalamic neurons that mirror aggression. Cell 186, 1195–1211 e1119 (2023). 10.1016/j.cell.2023.01.022

55 Kim, E. J., Jacobs, M. W., Ito-Cole, T. & Callaway, E. M. Improved Monosynaptic Neural Circuit Tracing Using Engineered Rabies Virus Glycoproteins. Cell Rep 15, 692–699 (2016). 10.1016/j.celrep.2016.03.067

56 Ciabatti, E., Gonzalez-Rueda, A., Mariotti, L., Morgese, F. & Tripodi, M. Life-Long Genetic and Functional Access to Neural Circuits Using Self-Inactivating Rabies Virus. Cell 170, 382–392 e314 (2017). 10.1016/j.cell.2017.06.014

57 Lim, N. K. et al. An Improved Method for Collection of Cerebrospinal Fluid from Anesthetized Mice. J Vis Exp (2018). 10.3791/56774

